# Mechanisms of generation of membrane potential resonance in a neuron with multiple resonant ionic currents

**DOI:** 10.1101/126714

**Authors:** David M. Fox, Hua-an Tseng, Tomasz G. Smolinski, Horacio G. Rotstein, Farzan Nadim

## Abstract

Neuronal membrane potential resonance (MPR) is associated with subthreshold and network oscillations. A number of voltage-gated ionic currents can contribute to the generation or amplification of MPR, but how the interaction of these currents with linear currents contributes to MPR is not well understood. We explored this in the pacemaker PD neurons of the crab pyloric network. The PD neuron MPR is sensitive to blockers of H- (*I*_H_) and calcium-currents (*I*_Ca_). We used the impedance profile of the biological PD neuron, measured in voltage clamp, to constrain parameter values of a conductance-based model using a genetic algorithm and obtained many optimal parameter combinations. Unlike most cases of MPR, in these optimal models, the values of resonant- (*f*_res_) and phasonant- (*f*_φ=0_) frequencies were almost identical. Taking advantage of this fact, we linked the peak phase of ionic currents to their amplitude, in order to provide a mechanistic explanation the dependence of MPR on the *I*_Ca_ gating variable time constants. Additionally, we found that distinct pairwise correlations between *I*_Ca_ parameters contributed to the maintenance of *f*_res_ and resonance power (Q_Z_). Measurements of the PD neuron MPR at more hyperpolarized voltages resulted in a reduction of *f*_res_ but no change in Q_Z_. Constraining the optimal models using these data unmasked a positive correlation between the maximal conductances of *I*_H_ and *I*_Ca_. Thus, although *I*_H_ is not necessary for MPR in this neuron type, it contributes indirectly by constraining the parameters of *I*_Ca_.

**Author Summary:** Many neuron types exhibit membrane potential resonance (MPR) in which the neuron produces the largest response to oscillatory input at some preferred (resonant) frequency and, in many systems, the network frequency is correlated with neuronal MPR. MPR is captured by a peak in the impedance vs. frequency curve (Z-profile), which is shaped by the dynamics of voltage-gated ionic currents. Although neuron types can express variable levels of ionic currents, they may have a stable resonant frequency. We used the PD neuron of the crab pyloric network to understand how MPR emerges from the interplay of the biophysical properties of multiple ionic currents, each capable of generating resonance. We show the contribution of an inactivating current at the resonant frequency in terms of interacting time constants. We measured the Z-profile of the PD neuron and explored possible combinations of model parameters that fit this experimentally measured profile. We found that the Z-profile constrains and defines correlations among parameters associated with ionic currents. Furthermore, the resonant frequency and amplitude are sensitive to different parameter sets and can be preserved by co-varying pairs of parameters along their correlation lines. Furthermore, although a resonant current may be present in a neuron, it may not directly contribute to MPR, but constrain the properties of other currents that generate MPR. Finally, constraining model parameters further to those that modify their MPR properties to changes in voltage range produces maximal conductance correlations.

## Introduction

Neuronal network oscillations at characteristic frequency bands emerge from the coordinated activity of the participating neurons. Membrane potential resonance (MPR) is defined as the ability of neurons to exhibit a peak in their voltage response to oscillatory current inputs at a preferred or resonant frequency (*f*_res_) [1]. MPR has been observed in many neuron types such as those in the hippocampus [2-4] and entorhinal cortex [2-6], inferior olive [7, 8], thalamus [1, 9], striatum [10, 11], as well as in invertebrate oscillatory networks such as the pyloric network of the crustacean stomatogastric ganglion (STG) [12-14]. Neurons may also exhibit phasonance or a zero-phase response, which describes their ability to synchronize with oscillatory inputs at a preferred phasonant frequency (*f*_φ=0_) [4, 15-18]. Resonance, phasonance and intrinsic oscillations are related, but are different phenomena as one or more of them may be present in the absence of the others [15, 16, 18].

Resonant and phasonant frequencies result from a combination of low- and high-pass filter mechanisms produced by the interplay of the neuron’s passive properties and one or more ionic currents and their interaction with the oscillatory inputs [1, 15, 18, 19]. The slow resonant currents (or currents having resonant gating variables) oppose voltage changes and act as high-pass filters. They include the hyperpolarization-activated inward current (*I*_H_) and the slow outward potassium current (*I*_M_). On the other hand, the fast amplifying currents (or currents having amplifying gating variables) favor voltage changes and can make MPR more pronounced. They include the persistent sodium current (*I*_NaP_) and the inward rectifying potassium (I_Kir_) current. Most previous systematic mechanistic studies have primarily examined models with one resonant and one amplifying current, such as *I_H_* and *I*_NaP_, respectively [15, 18-20]. Currents having both activating and inactivating gating variables (in a multiplicative way) such as the low-threshold calcium current (*I*_Ca_) are not included in this classification, but they are able to produce resonance by mechanisms that are less understood [16, 21].

Although a causal relationship between the properties of MPR and network activity has not been established [but see 22], resonant neurons have been implicated in the generation of network oscillations in a given frequency band because the resonant and network frequencies often match up or are correlated. One example is in the hippocampal theta oscillations [23] in which CA1 pyramidal cells exhibit MPR *in vitro* at theta frequencies of 4-10 Hz [2-4, 24] (but see [25]). Interestingly, MPR is not constant across the somatodendritic arbor in these neurons [26]. Hippocampal interneurons also show MPR *in vitro*, but at gamma frequencies of 30-50 Hz [3, but see 4], and gamma oscillations have been found to be particularly robust in network models containing resonant interneurons [27, 28].

The crab pyloric network produces stable oscillations at a frequency of ^∼^1 Hz, driven by a pacemaker group composed of two neuron types, the anterior burster (AB) and the pyloric dilator (PD), that produce synchronized bursting oscillations through strong electrical-coupling [29]. The PD neuron shows MPR, with *f*_res_ ^∼^1 Hz that is positively correlated with the pyloric network frequency [12]. Previous work has demonstrated that MPR in this neuron depends on two voltage-gated currents: *I*_Ca_ and *I*_H_ [12]. Ionic current levels in pyloric neurons are highly variable across animals, even in the same cell type [30]. It is therefore unclear how these currents may interact to produce a stable MPR in the PD neuron and whether this variability persists or is increased or decreased in the presence of oscillatory inputs.

Traditionally, MPR is measured by applying ZAP current injection and recording the amplitude of the voltage response [1, 31]. In some systems, depolarization can increase [32] or decrease [33], 1996) the preferred frequency. Alternatively, resonance is measured by applying ZAP voltage inputs in voltage clamp and recording the amplitude of the total current. Both approaches yield identical results for linear systems, but not necessarily for nonlinear systems. A previous study from our lab using the voltage clamp technique showed that in the PD neuron hyperpolarization decreases both *f*_res_ and network frequencies [14]. Since MPR results from the outcome of the dynamics of voltage-gated ionic currents activated in different voltage ranges, changing the input voltage amplitude is expected to change *f*_res_ in an input amplitude-dependent manner. This cannot be captured by linear models in which impedance is independent of the input amplitude. To our knowledge, no study has attempted to understand the ionic mechanisms that produce shifts in *f*_res_ in response to changes in the voltage range.

Previous studies have explored the generation of MPR by *I*_Ca_ and through the interaction between *I*_Ca_ and *I*_H_ in hippocampal CA1 pyramidal neurons [16, 17] and thalamic neurons [21], where the resonant and network frequencies are significantly higher than in the crab pyloric network and the *I*_Ca_ time constants are smaller. Based on numerical simulations, these investigations have produced important results about the role of the activating and inactivating gating variables and their respective time constants play in the generation of MPR and the determination of *f*_res_. However, a mechanistic understanding of the effects of the interacting time constants and voltage-dependent inactivation that goes beyond simulations is lacking. An important finding for the CA1 pyramidal neurons is that, for physiological time constants, they exhibit resonance, but no phasonance [16]. However, for larger time constants, outside the physiological range for these neurons, they are able to exhibit phasonance. This suggests that PD neurons, which have slower time scale currents, may exhibit resonance and phasonance at comparable frequencies. If so, such a correlation between resonance and phasonance can be used to explain the influence of ionic current parameters.

Our study has two interconnected goals: (i) to understand how the interplay of multiple resonant gating variables shapes the Z- and ϕ-profiles (impedance amplitude and phase-shift as a function of input frequency) of a biological PD neuron, and (ii) to understand the many ways in which these interactions can occur to produce the same Z-profile in these neurons. For a neuron behaving linearly, e.g., with small subthreshold inputs, this task is somewhat simplified by the fact that linear components are additive. However, neurons are nonlinear and the nonlinear interaction between ionic currents has been shown to produce unexpected results [16, 18, 19].

To achieve these goals we measured and quantified the Z- and φ-profiles of the PD neuron. We then used a single-compartment conductance-based model of Hodgkin-Huxley type [34] that included a passive leak and the two voltage-gated currents *I*_H_ and *I*_Ca_ to explore what combinations of model parameters can produce the experimentally observed PD neuron Z- and φ-profiles. The maximal conductances of ionic currents of neurons in the stomatogastric nervous system vary widely [35-37]. We therefore assume that the parameters that determine the Z-profile in the PD neuron vary across animals. Thus, instead of searching for a single model that fit the PD neuron Z-profile, we used a genetic algorithm to capture a collection of parameter sets that fit this Z-profile. To achieve such a fit, we defined a set of ten attributes that characterize the PD neuron Z-profile (e.g., resonant frequency and amplitude) and used a multi-objective evolutionary algorithm [MOEA, 38] to obtain a family of models that fit these attributes. We then used this family of optimal models to identify the important biophysical parameters and relationships among these parameters to explain how the PD neuron Z-profile is shaped. We show how the fact that the inactivating calcium current peaks at the same phase as the passive properties, in response to sinusoidal inputs, can explain why resonant and phasonant frequencies are equal. We identify significant pairwise parameter-correlations, which selectively set certain attributes of MPR. We show that, in this neuron, *I*_H_ does not produce MPR but can extend the dynamic range of *I*_Ca_ parameters mediating MPR. Furthermore, we identify a subset of models that capture the experimental shift in the resonant frequency with changes in lower bound of voltage oscillation. Finally, we exploit the fact that the resonant and phasonant frequencies are equal for the PD neuron to provide a mechanistic understanding of the effects of the *I*_Ca_ time constants on the resonant frequency by using phase information. Our results provide a mechanistic understanding for a generic class of neurons that exhibit both resonance and phasonance as the result of the interaction between multiplicative gating variables and complement the studies in [16].

## Results

The PD neuron produces 1 Hz bursting oscillations with a slow-wave approximately -60mV to -30mV (fig 3a). Driving the neuron through this voltage range with a ZAP function in voltage clamp (fig 3b top panel) produces a minimum (arrow in fig 3b bottom panel) in the amplitude of the current response (fig 3b). The input frequency at which this minimum occurs corresponds to a peak in the Z-profile (*f*_res_, *Z*_max_; fig 3c1). The value of *f*_res_ was 0.86 ± 0.05Hz producing *Z*_max_ values of 10.23 ± 0.51 MΩ (N = 18; fig 3d). The *φ*-profile shows a phasonant frequency *f_φ_*_=_*_0_* = 0.81 ± 0.05Hz, which in most cases matched *f*_res_ (fig 3c2). The PD neuron had a *Q_Z_* of 2.77 ± 0.71 MΩ and Λ_ν_ of 0.53 ± 0.04 Hz. Across preparations, *Q_Z_* showed considerable variability, whereas *f*_res_, Λ_½_, and *f_φ_*_=_*_0_* were relatively consistent (fig 3d). The corresponding median values for *f*_res_, *Q_Z_*, Λ_½_, and *f_φ_*_=_*_0_* were 0.83 Hz, 2.77 MΩ, 0.5 Hz, 0.79 Hz, respectively.

**Fig 1.**
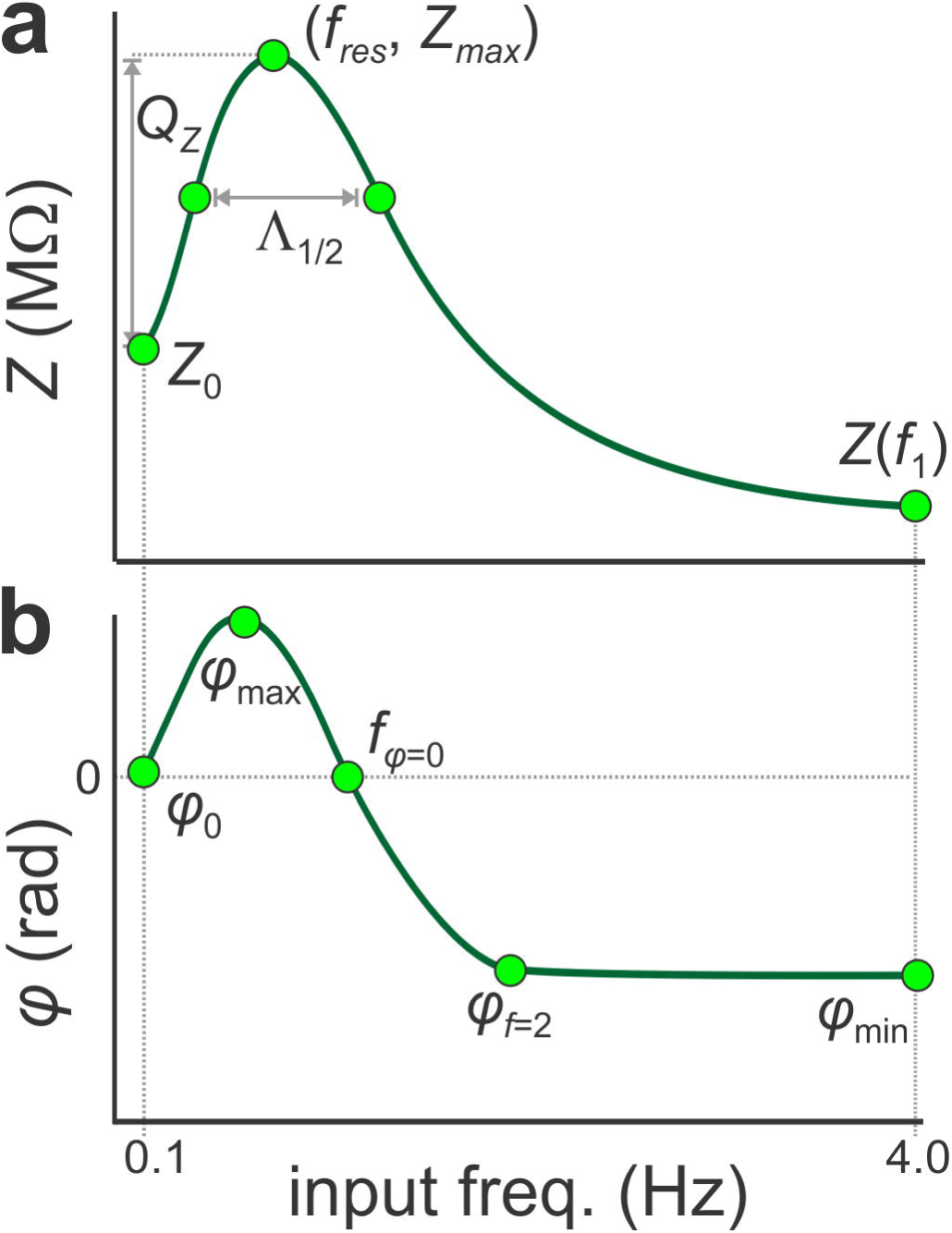
Characterization of impedance amplitude *Z(f)* and phase *φ(f)* into target objective functions was performed to constrain the model parameters. The individual objective functions which collectively measure goodness-of-fit were taken as the distance away from characteristic points along the *Z*(*f*) and *φ*(*f*) profiles (green circles). **a.** The attributes used along *Z*(*f*) were *Z*_0_=*Z*(*f*_0_) at *f*_0_=0.1 Hz, *Z*(*f_1_*) atf_1_ = 4 Hz, maximum impedance *Z*_max_=(Z*f*_res_) and the two points of the profile at *Z*_0_+*Q_Z_*/2. *Q*_Z_=*Z*_max_-*Z*_0_. Λ_½_ is the width of the profile at *Z*_0_+*Q_Z_*/2. **b.** The attributes used along *Z*(*f*) were *Z*(*f*_0_), maximum advance *φ*_max_, zero-phase frequency *f*_φ=0_, φ*_f_*_=2_ at 2 Hz and maximum delay *φ*_min_.

**Fig 2.**
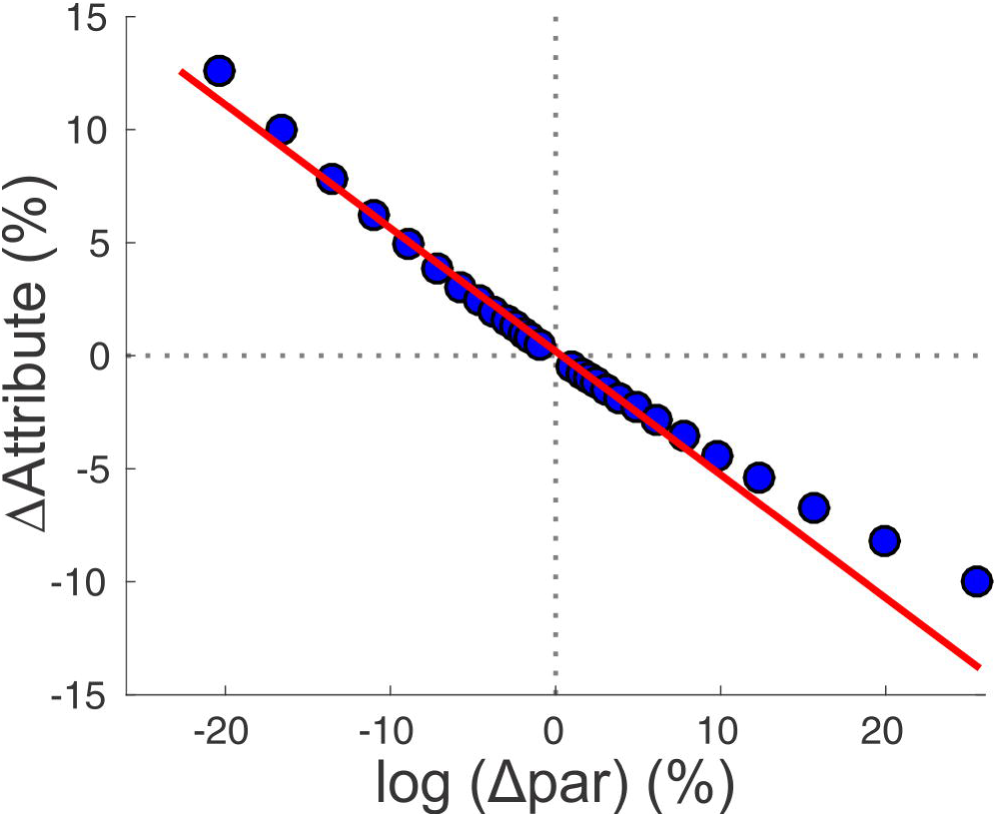
Linear fits used to assess the sensitivity of impedance attributes on changes in parameters. Each model parameter was changed from the optimal value (origin) in both directions on a logarithmic scale to characterize parameter sensitivity. The slope of a linear fit of the relative change in the *Z*(*f*) attribute and the parameter was measured as sensitivity. The parameter was changed until the fit was no longer linear (R^2^<0.98).

**Fig 3.**
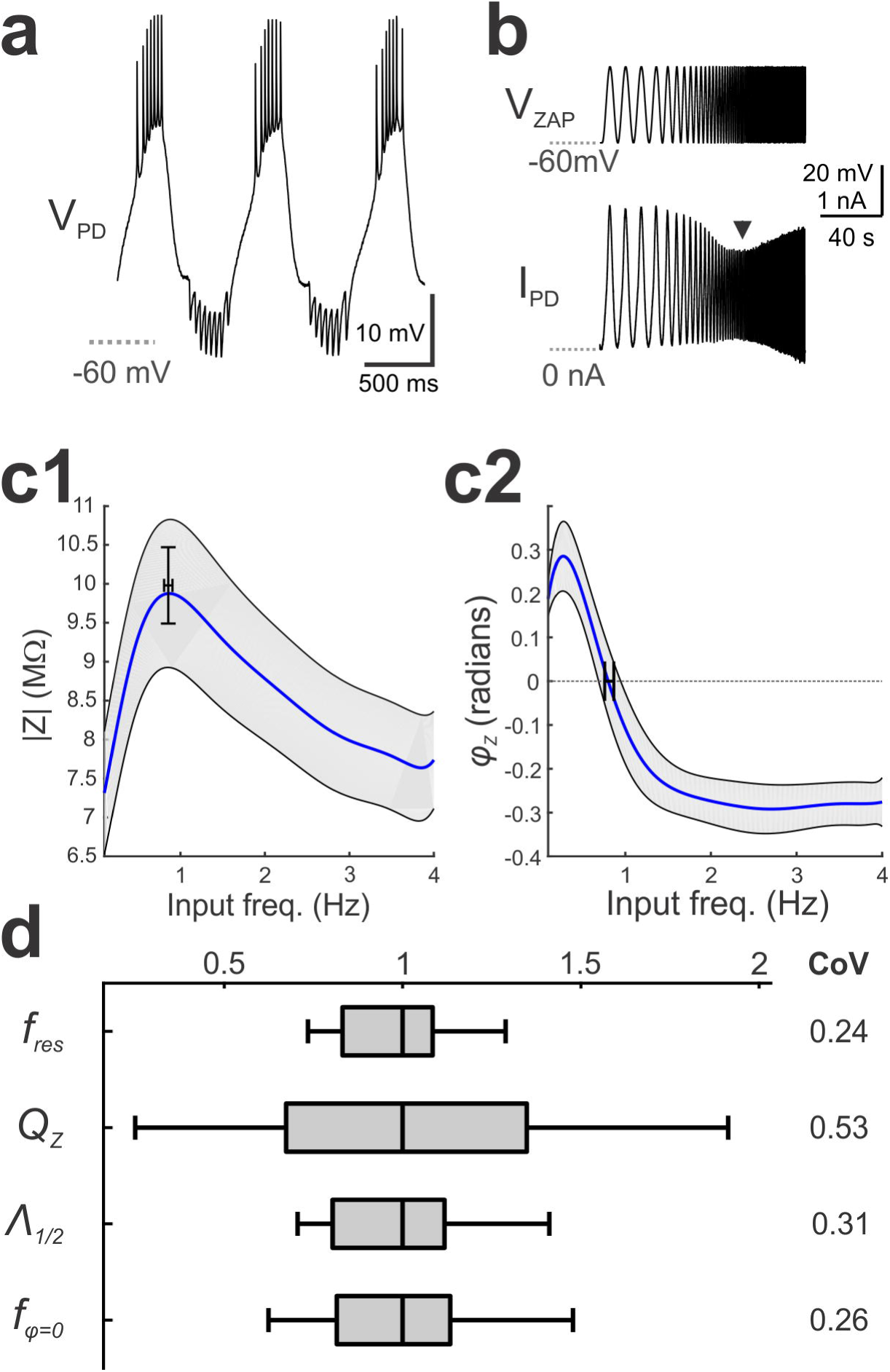
Membrane potential resonance MPR of the PD neuron was measured in voltage clamp. **a.** During ongoing activity, the PD neuron shows a slow-wave voltage waveform ranging approximately between - 60 and -30 mV. **b**. The membrane potential (V_zap_) and the injected current (I_PD_) were recorded when the PD neuron was voltage-clamped using a ZAP function between -60 and -30mV and sweeping frequencies between 0.1 and 4 Hz. The arrowhead indicates resonance, where the current amplitude is minimal and Z is maximal. **c.** The impedance amplitude *Z*(*f*) (**c1**) and phase *φ*(*f*) (**c2**) profiles of the PD neuron recorded in 18 preparations. The cross bars show the mean and SEM of *f*_res_ and *Z*_max_ (**c1**) and *f_v=0_* (**c2**). The shaded region indicates the 95% confidence interval. d. The range of three *Z*(*f*) attributes *f*_res_, *Q*_Z_, and *φ_1_/_2_* and one *φ*(*f*) attribute *f*_*φ*=0_. Each attribute was normalized to the median of its distribution for cross comparison. CoV is the coefficient of variation.

To obtain model parameter combinations constrained by the PD neuron *Z*- and ϕ - profiles, we generated a population of models using an NSGA-II algorithm (see Methods). The attributes of a single PD neuron *Z*- and ϕ -profiles (fig 4, filled red circles) constrained the optimization of the parameter values. This resulted in a population of ^∼^9000 sets of parameters (“optimal” dataset). All models in the optimal dataset captured the attributes of *Z* and φ to within 5% of the target (light blue lines in fig 4), with the exception of *φ*_max_, which may be due to the anatomical structure of the PD neuron, a property that is omitted in our single-compartment model, or due to additional ionic currents, such as the potassium A current, which are not included in our model [16, 39].

**Fig. 4.**
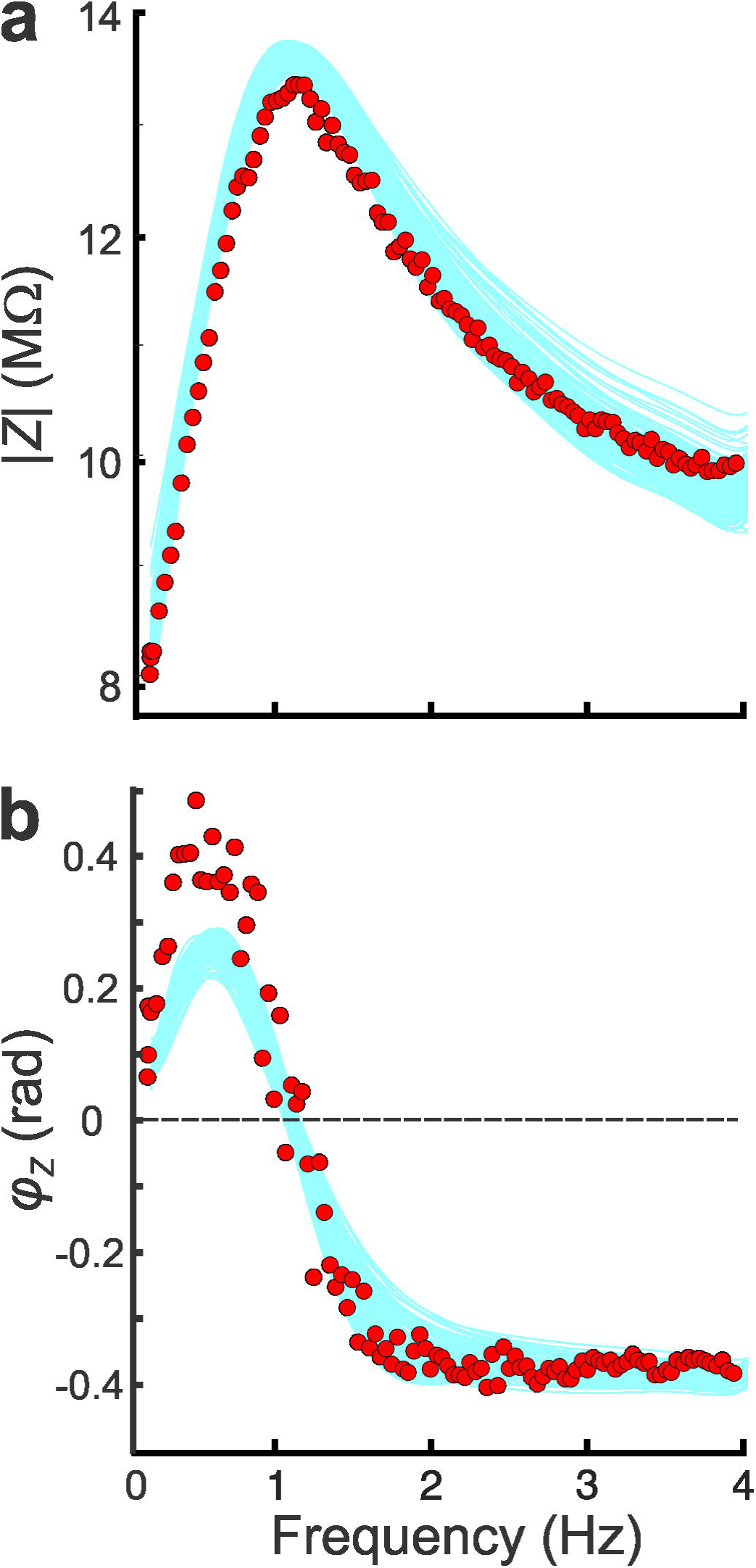
Optimal models were fit to the impedance attributes of a single PD neuron. The *Z*(*f*) (**a**) and *φ*(*f*) (**b**) profiles of 500 randomly selected models from the optimal dataset (light blue curves) are compared to the target neuron’s impedance profiles (red circles). All attributes (except *φ*_max_) were captured to within 5% accuracy. The values of the biological target impedance amplitude attributes (in Hz, MΩ) were: *f*_0_, Z_0_) = (0.1, 8.2), *f*_res_, *Z*_max_) = (1, 13.7), (0.4, 11.65), (2.5, 11.65) and (4, 9.6). The target impedance phase attributes (in Hz, rad) were: (0.1, 0), *f*_*φ*max_, *φ*_max_) = (0.4, 0.5), *f*_*φ*=0_, 0) = (1.05, 0), (2, -4), (*f_φ_*_min_, *φ*_min_) = (4, -0.4).

### The generation of MPR by the interaction of two resonant voltage-gated currents

To understand how *Z* is generated by the dynamics of individual ionic currents at different voltages and frequencies, we examined the amplitude and kinetics of ionic currents. In voltage clamp, *Z* is shaped by active voltage-gated currents, interacting with the passive leak and capacitive currents, in response to the voltage inputs. To understand the contribution of different ionic currents, we measured these currents in response to a constant frequency sine wave voltage inputs (fig 5a inset) at three frequencies: 0.1Hz, 1Hz (*f*_res_) and 4Hz (fig 5). For these frequencies, we plotted the steady-state current as a function of voltage (fig 5b-d left) and normalized time (or cycle phase = time x frequency; fig 5b-d right). At 0.1 Hz, the amplitudes of *I*_H_ and *I*_L_ +*I*_Cm_ sets *I*_total_ at low (^∼^ -60 mV) and high (^∼^ -30 mV) voltages, respectively (fig 5b left). Since *I*_H_ deactivation is slow, it also contributes to *I*_total_ at high voltages (fig 5b right). At 1 Hz (= *f*_res_), *I*_H_ still sets the minimum of the total current, but, because of its slow kinetics, its steady-state dynamics are mostly linear (fig 5c left). However, now *I*_Ca_ peaks in phase (fig 5c right) with the passive *I*_L_ + *I*_Cm_ at high voltages, thus producing a smaller *I*_total_ (magenta bar in fig 5c). The values of *I*_H_ at 4 Hz are not much different from 1 Hz (fig 5d). However, *I*_Ca_ peaks at a much later phase (fig 5d right), which does not allow it to compensate for *I*_L_ + *I*_Cm_ at high voltages, thus resulting in a larger *I*_total_ (magenta bar in fig 5d). Note that at 1 Hz, the total current peaks at a cycle phase close to 0.5, thus implying that that the *f*_res_ and *f*_φ=0_ are very close or equal (fig 5c right). Although figure 5 shows the results for only one model in the optimal dataset, these results remain nearly identical for all models in the optimal dataset. The standard deviation of the currents measured, including the total current was never above 0.15 nA over all models. The inset in fig. 5c shows one standard deviation around the mean for the data shown in the right panel, calculated for 200 randomly selected models.

**Fig 5.**
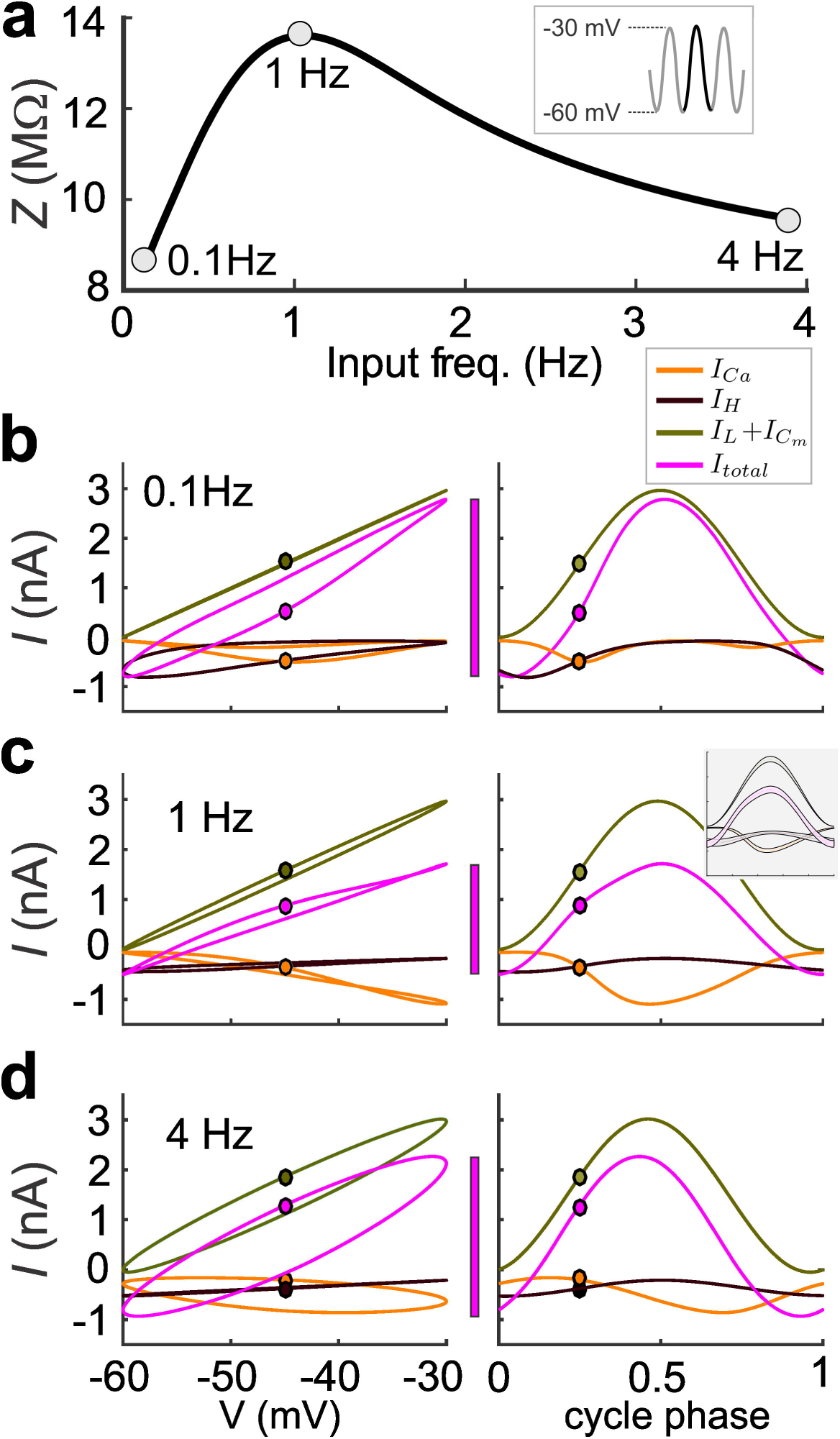
Passive and voltage-gated currents contribute to the generation of MPR. **a.** *Z*(*f*) for a random model from the optimal dataset. We measured the steady-state response to sinusoidal voltage inputs (inset) at 0.1 Hz, *f*_res_=1 Hz, and 4 Hz. Voltage-gated (*I*_Ca_ and I_H_) and passive currents (*I*_L_ + *I*_Cm_) are plotted as a function of voltage (left) and normalized time or cycle phase (right) at 0.1 Hz **(b),** 1 Hz **(c)**, and 4Hz **(d).** The inset in **5c** shows one standard deviation around the mean for the data shown in the right panel, calculated for 200 randomly selected models.

An important collective property of the models we found is that the two frequencies, *f*_res_ and *f*_φ=0_ coincide (fig. 6a-b). We analyzed the experimental data, and confirmed that the coincidence of MPR and phasonance frequencies also occurs in the biological system (fig. 6b inset). This is typically not the case for neuronal models (and for dynamical systems in general), not even for linear systems [18-20], with the exception of the harmonic oscillator. However, the fact that it occurs in this system, allows us to use the current vs. cycle phase (current-phase) diagrams to understand the dependence of *f*_res_ and *f*_φ=0_ on the model parameters (fig. 6c).

**Fig 6.**
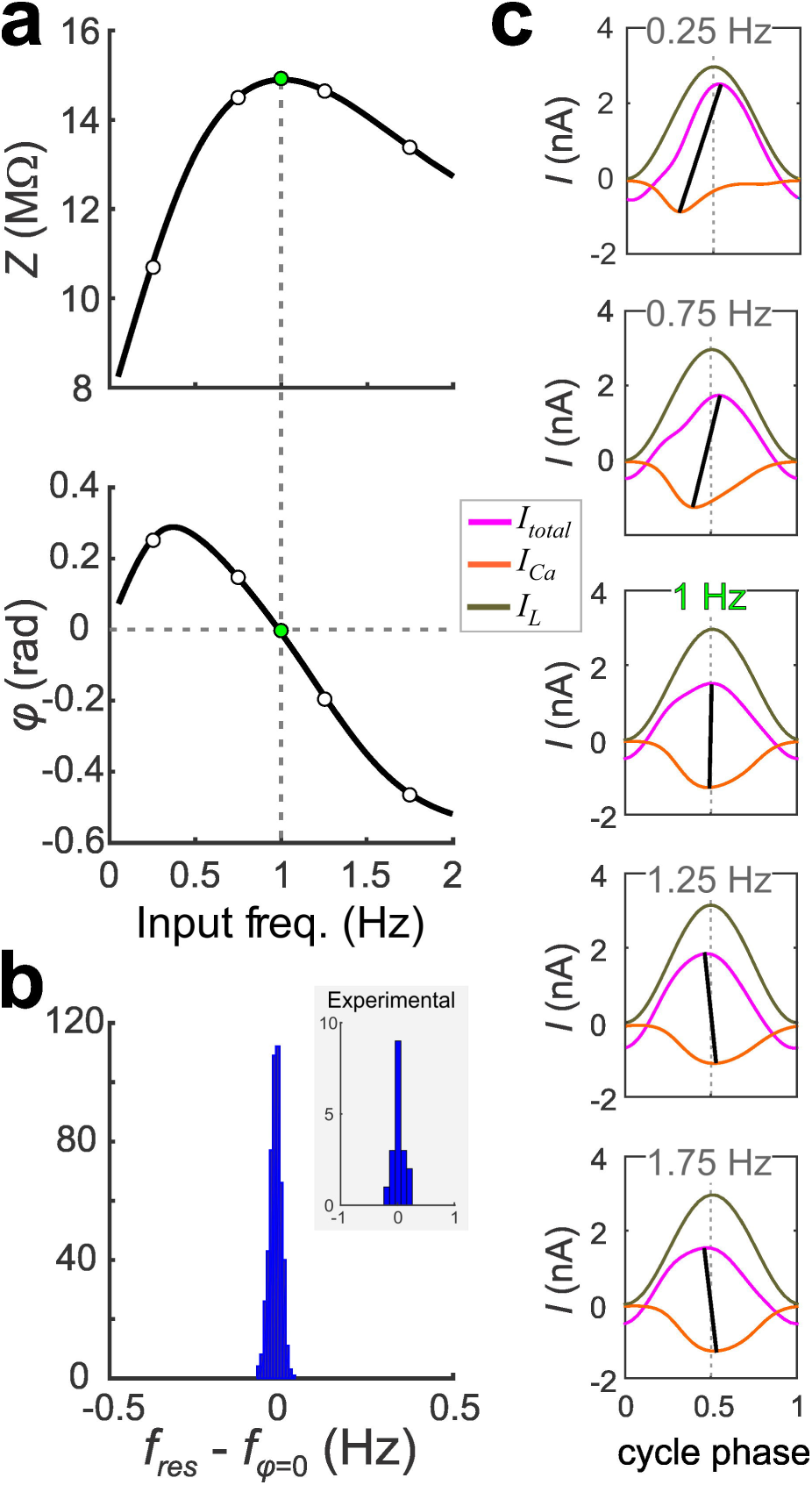
*f*_res_ and *f*_φ=0_ of the optimal models are nearly identical. **a.** *Z*(*f*) (top) and *φ*(*f*) (bottom) for a representative optimal model. Green dots indicate *f*_res_ (top) and *f*_φ=0_ (bottom). **b**. Histogram showing the difference between *f*_res_ and *f*_φ=0_ for 500 randomly selected models. A comparison of *f_r_*_es_ and *f*_φ=0_ of the experimental data of the PD neuron shows a similar distribution (inset, N=18). (**c**) Plots of steady-state responses of *I*_Ca_, *I*_L_, and *I*_total_ to sinusoidal voltage inputs at the frequencies marked in panel **a** shown as a function of normalized time (cycle phase). Dotted vertical line indicates cycle phase 0.5 where the passive currents peak. Solid lines connect the minimum of *I*_Ca_ to the peak of *I*_total_. The two lines nearly align at *f*_φ=0_.

The current-phase diagrams are depicted as in fig 5b-d, as graphs of *I*_total_, *I*_L_ and *I*_Ca_ as a function of the cycle phase for each given specific input frequency (fig. 6c). We do not show *I*_H_ and *I*_Cm_ in this plot, because at frequencies near *f*_res_ they do not change much with input frequency. Note that *I*_L_ is independent of the input frequency (five panels in fig. 6c) because it precisely tracks the input voltage.

In voltage clamp, *f*_φ=0_ = 1Hz is where I_total_ is at its minimum amplitude exactly at cycle phase 0.5, coinciding with the peak of the input voltage (fig. 6c, middle). The fact that *I*_L_ precisely tracks the input voltage imposes a constraint on the shapes of *I*_Ca_ and *I*_total_. Therefore, by necessity, if the *I*_Ca_ trough occurs for a cycle phase below 0.5, the *I*_total_ peak must occur for a cycle phase above 0.5 (fig. 6-c, top two panels) and vice versa (fig. 6c, bottom two panels). This is shown by the slope of the line joining the peaks of *I*_total_ and *I*_Ca_ and, at *f*_res_ this line is approximately vertical (fig. 5c middle panel).

We use this tool to explain the dependence of the *Z*-profile on the time constants 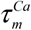 (fig. 7a) and 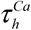 (fig. 7b). The corresponding current-phase diagrams are presented in figs. 7c and 7d, respectively. In each panel we present the current-phase diagrams for *f* at 1 Hz (=*f*_res_ when the parameter is at 100%; middle) and *f* = *f*_res_ (sides) when *f*_res_ is different from 1Hz.

**Fig 7.**
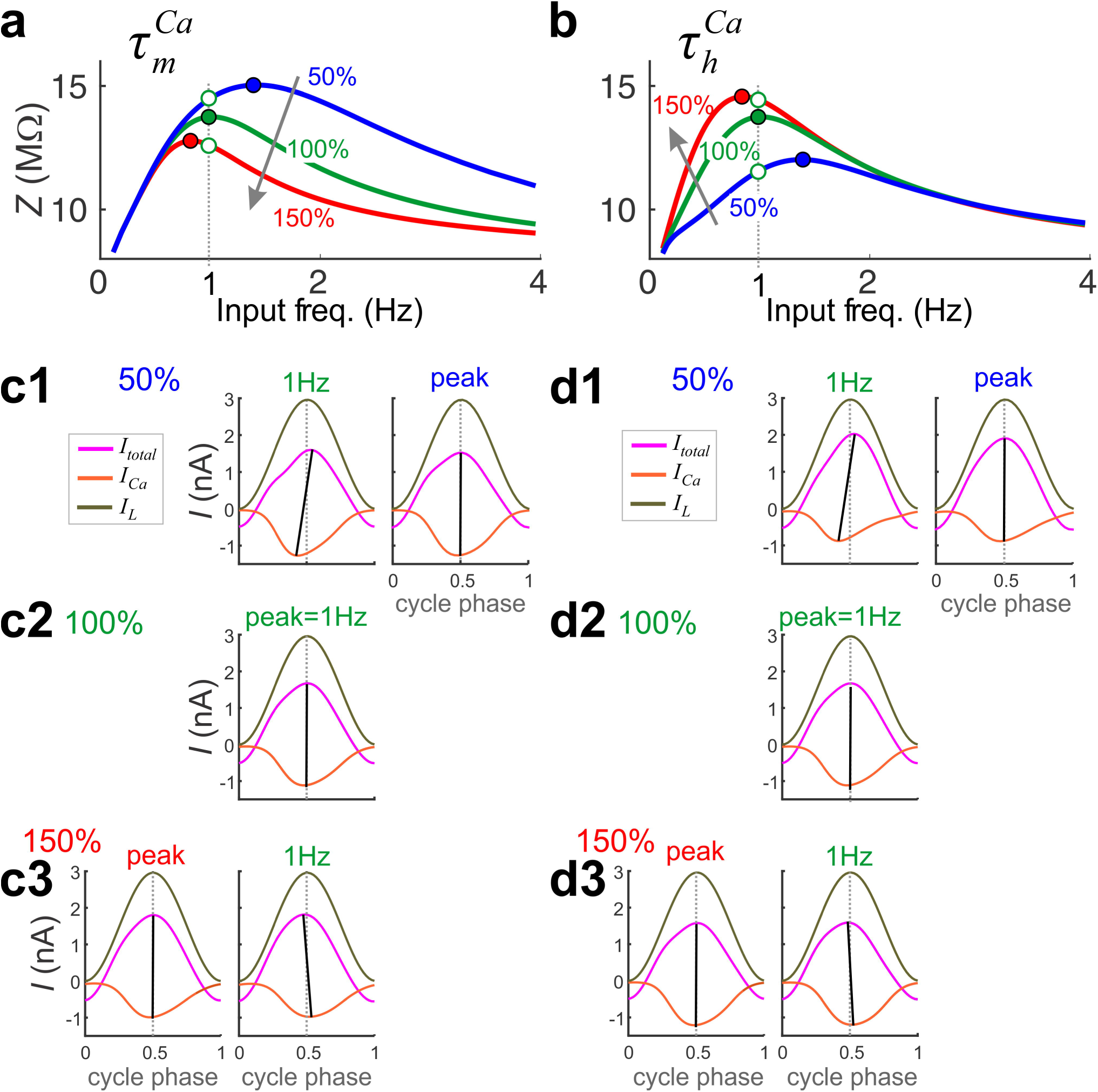
The time constants of I_Ca_ activation and inactivation control *f*_res_ and *Z*_max_. The *Z*(*f*) profiles are plotted for a randomly selected optimal model (green) at different values of 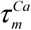 (**a**) and 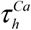 (**b**). Note that *f*_res_ of the control (100%) values are at 1 Hz (dashed vertical line). The currents *I*_Ca_, *I*_L_ and *I*_total_ plotted as a function of cycle phase at 50% (**c1, d1**), 100% (**c2, d2**), and 150% (**c3, d3**) of the control values of 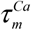 and 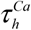 (**d**). In each panel of c and d, the currents are shown at 1 Hz (along the dashed lines in **a, b**) and at *f*_res_ (filled circles in **a, b**).

To understand the dependence of *Z* on changes in 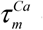 and 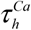 we have to primarily explain the dependence of the two attributes *Z*_max_ and *f*_res_ on these parameters. While *f*_res_ has a similar monotonic dependence on 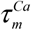 and 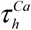 (as these parameters increase, *f*_res_ decreases), *Z*_max_ has the opposite dependence on 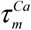 and 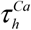. The opposite dependence of *Z*_max_ on 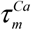 and 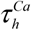 is a straightforward consequence of the opposite feedback effects (positive for 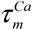 and negative for 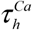) that these parameters exert on *I*_Ca_. An increase in 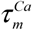 (for fixed values of 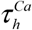) results in a smaller *I*_Ca_ in response to a given voltage clamp input. Because *I*_Ca_ is smaller and negative, this leads to an increase in *I*_total_ and a decrease in *Z* at all frequencies. Similarly, an increase in 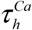 (for fixed values of 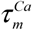) results in a larger *I*_Ca_, leading to a decrease in *I*_total_ and an increase in *Z*.

For a fixed value of the input frequency *f* (e.g. *f* = 1 Hz as in fig. 7), for *Z*_max_ to decrease as 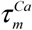 increases (fig. 7-a), the cycle phase of peak *I*_Ca_ is delayed thereby subtracting less from *I*_L_ on the depolarizing phase. This leads to *I*_total_ to phase advance relative to *I*_L_ (fig. 7-c) and causes *f*_res_ to decrease. Similarly, for *Z*_max_ to increase as 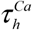 increases (fig. 7-b), *I*_Ca_ has to peak later in the cycle thereby subtracting less from *I*_L_ on the depolarizing phase, which causes *I*_total_ to peak earlier in the cycle, which in turn causes the bar also to swing from the left to the right (fig. 7-d). Therefore, *f*_res_ decreases.

### Parameter constraints and pairwise correlations

Previous studies have shown that stable network output can be produced by widely variable ion channel and synaptic parameters [37, 40]. Our biological data, similarly, showed that many of the Z- and ϕ- profile attributes, such as *f*_res_, Λ _½_ and *f*_*φ*=0_ are relatively stable across different PD neurons whereas *Q_Z_* shows the most variability (fig 3d). To determine whether the Z- and ϕ-profile attributes constrain ionic current parameters, we examined the variability of the model parameters in the optimal dataset. We found that some parameters were more constrained while others were widely variable, as measured by the coefficient of variation (CoV; fig 8a). Parameters showing large CoVs were *ḡ_Ca_*, 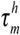, *ḡ_h_*, 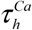, and 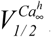; those showing small CoVs were *ḡ_L_* and the time constant of activation of I_H_ and I_Ca_ and half-activation voltage of *l_Ca_:* 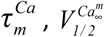, *ḡ_L_* (in increasing order of CoV value). A small CoV value implies that the parameter is tightly constrained in order to produce the proper Z- and *φ*-profiles.

**Fig 8.**
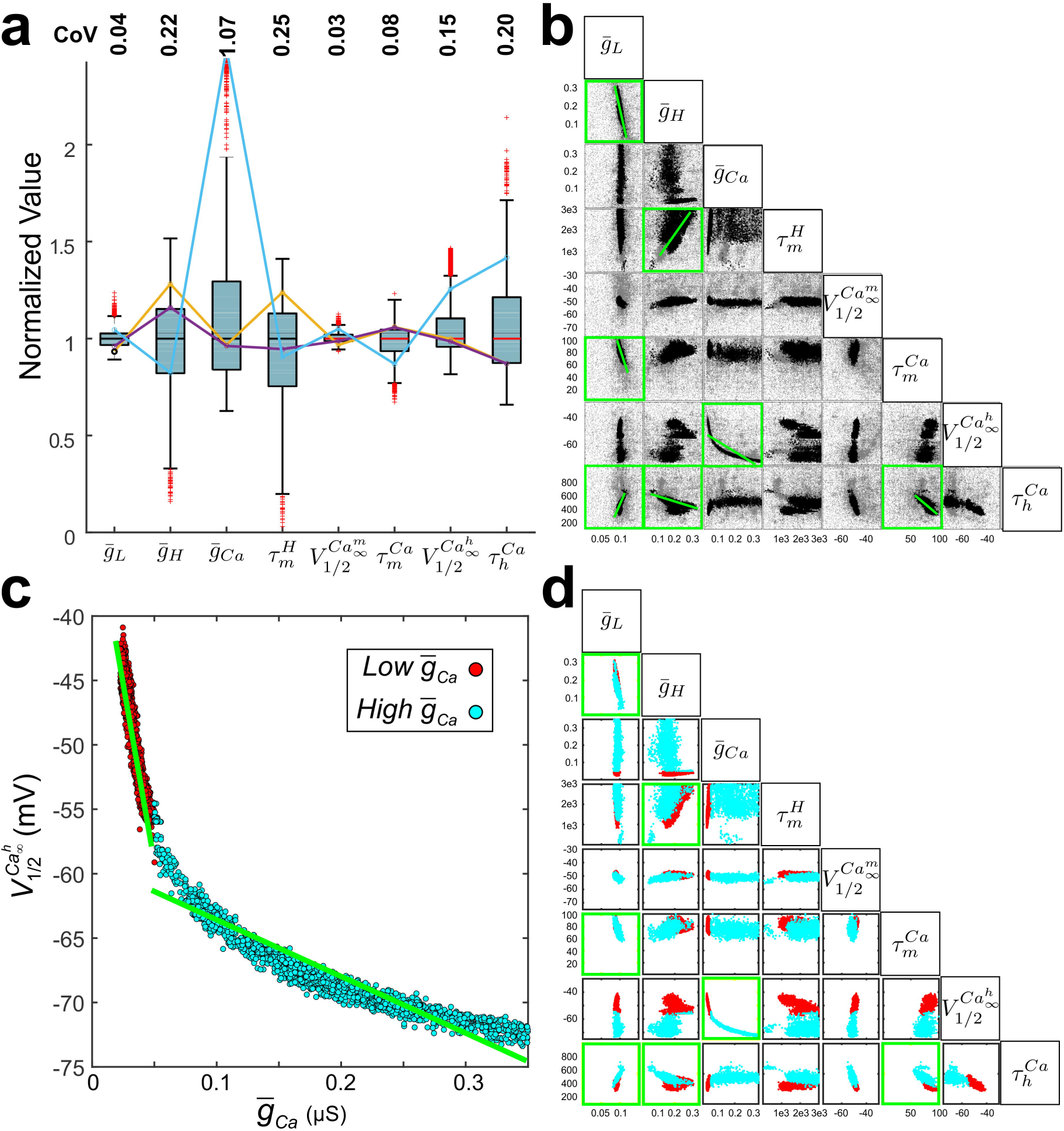
The optimal models show variability in individual and pairs of parameters. **a**. The range of parameters for all optimal models (^∼^9000). Each parameter is normalized by its median value for cross comparison. The median values were *ḡ_L_* = 0.096 *μS*, *ḡ_H_* = 0.164 *μS*, *ḡ_ca_* = 0.172 *μS*, 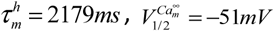, 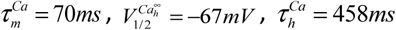 Three representative optimal model parameter sets are shown (cyan, orange, purple solid line segments) indicating that widely different parameter combinations can produce the biological *Z*(*f*) and φ(*f*). CoV is coefficient of variation. **b.** Pairwise relationships among parameters of all optimal models (black dots). The range of parameter space was sampled within the prescribed limits given to the optimization routine, shown by including the sampled non-optimal models (grey). Permutation test showed significant pairwise correlations (green highlighted boxes with linear fits shown as green lines). c. Optimal models could be separated into two highly significant linear fits (green lines) in *ḡ_ca_* 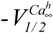 according to whether *ḡ_ca_* < 0.05 (red;Low *ḡ_ca_*) or *ḡ_ca_* >0.05 (cyan; High *ḡ_ca_*). **d.** All pairwise relationships, separated on the low or high *ḡ_ca_* (colors as in panel **c**). Green boxes are the same as in **b**.

A number of studies have indicated that the large variability in ion channel parameters is counter-balanced by paired linear covariation of these parameters [36, 37, 41-43]. Considering the large variability, we identified parameter pairs that co-varied (fig 8b). For this, we carried out a permutation test for the Pearson’s correlation coefficients, followed by a Student’s t-test on the regression slopes, to identify significant correlations between pairs of parameters (see Methods). The strongest correlations were between the following parameters: *ḡ_L_* - *ḡ_H_* (r=-0.93), *ḡ_L_-* 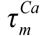 (R = 0.73), *ḡ_L_* -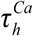 (R = 0.88), *ḡ_H_ -*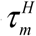 (R = 0.68),*ḡ_H_* -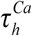 (R = -0.82), *ḡ_H_ -* 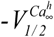 (R = 0.76), *ḡ_ca_* -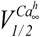 (R = -0.94), and 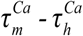 (R = -0.80) (correlations selected with p < 0.01; fig 8b).

In our experiments, 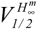 was fixed at -70 mV, using data from experimental measurements in crab [44] (see Methods). However, we also repeated the MOEA with 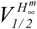 set to -96 mV, as reported in lobster experiments [45], and found that all correlations observed with the former value of 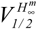 remain intact, but simply with a much larger maximal conductance of *I*_H_ (fig. S1). In other words, shifting 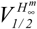 to the left simply results in larger *ḡ_H_* in the optimal models without qualitatively changing the other findings.

In particular, we found that the *ḡ_ca_ -* 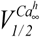 correlation appeared nonlinear, but there were strong and distinct linear correlations in the two regions the *ḡ_ca_* > 0.05 µS (low *ḡ_ca_*) and *ḡ_ca_* < 0.05 µS (high *ḡ_ca_*; fig 8c). To ensure that our partitioning of the population into different levels of *ḡ_ca_* was valid, we ran the MOEA two additional times, each time using only the mean values of *ḡ_L_*, 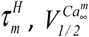, and 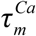 for either the low or the high *ḡ_ca_* values. These optimal models consistently separated into two non-overlapping model parameters, consistent with the low and high *ḡ_ca_* models in fig 8c.

We examined if the low and high *ḡ_ca_* models separated or showed distinct correlations in the remaining parameters. The two groups produced non-overlapping subsets of model parameters in the ḡ_ca -_ 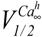 graph. We calculated the Pearson’s correlation coefficient for each pair of parameters in the low and high *ḡ_ca_* groups and tested for significance as before (see Table 2). We found that only the high *ḡ_ca_* group showed a significant 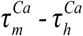 and *ḡ_ca_* −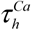 correlations (Table 2). Additionally, both low and high *ḡ_ca_* groups showed the following correlations: 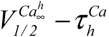, *ḡ_L_* - *ḡ_H_*, ḡ_ca -_ 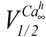 and ḡ_H-_ 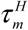, ḡ_ca -_ 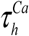Furthermore, when we ran the MOEA on models where *ḡ_H_* was set to 0, the only optimal models obtained fell within a narrow range of the high *ḡ_ca_* group (fig S2), which is consistent with the distribution of high *ḡ_ca_* models in the *ḡ_H_* − *ḡ_ca_* panel of figure 8d.

**Table 1.**
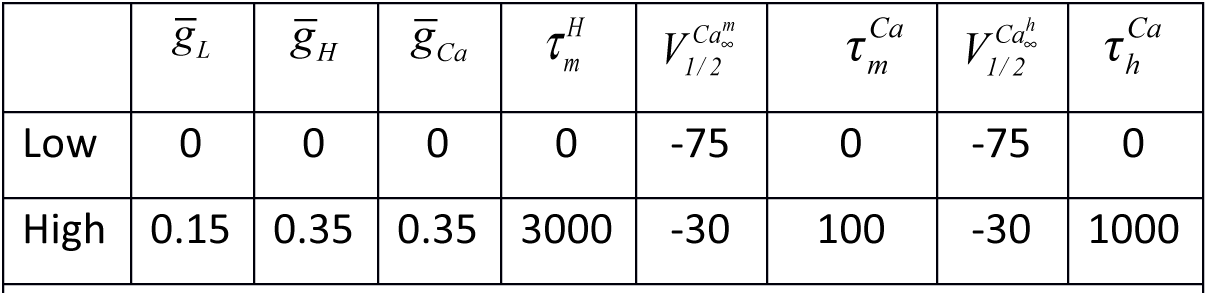
Limits of parameter values allowed for the PD neuron models. 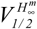 was fixed at -70 mV since there is little variability in the reporting of this experimental measurement [45, 61]. Voltages are in mV, maximal conductances in μS and time constants in ms.

**Table 2.**
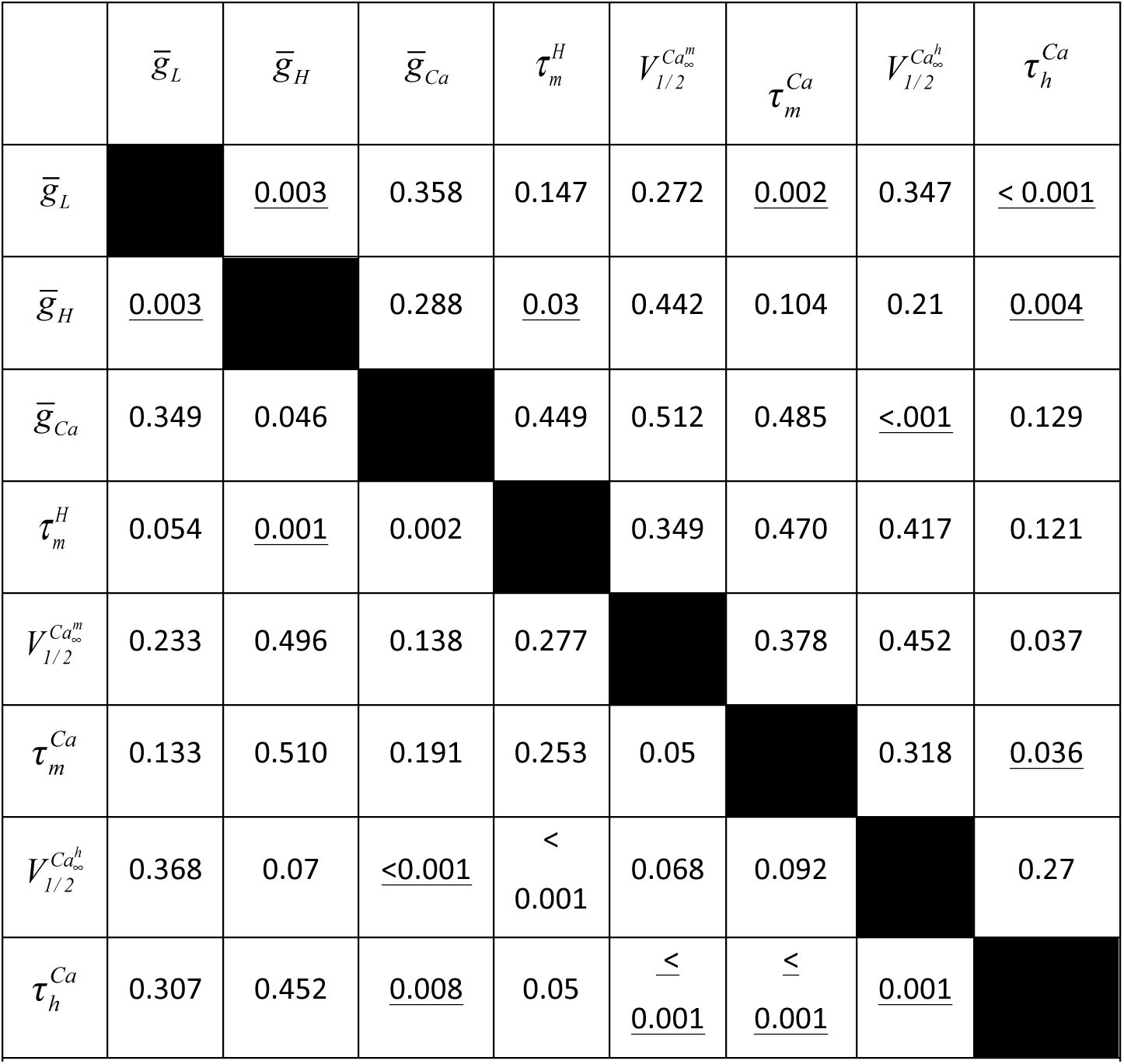
Statistical p-values obtained using the permutation test of pairwise comparisons for low (lower triangle) and high (upper triangle) *ḡ_ca_*. Underlined values are statistically significant (p<0.05).

### Decreasing the lower bound of voltage oscillations influences the measured *f*_res_ and *Z*_max_

The lower voltage range of the PD bursting oscillation is strongly influenced by the inhibitory synaptic input from the lateral pyloric neuron (LP), and previous work has shown that *f_res_* in the PD neuron is influenced by the minimum of the voltage oscillation (*V*_low_) [14]. In order to explore which subset of our optimal models faithfully reproduce the influence of the minimum voltage range, we measured the Z-profile when *V*_low_ was changed from - 60 to -70 mV (fig 9a). Decreasing V_low_ significantly decreased *f*_res_ (by 0.24±0.8Hz), while there was no significant difference in the mean *Z*_max_ (-0.15±0.81MΩ) (two-way RM-ANOVA; N = 8, p < 0.001; fig 9b, left panel).

To explore whether the shift in *f*_res_ as a function of *V*_low_ could be captured by either low or high *ḡ_ca_* models, we measured the shift in *f*_res_ and *Z*_max_, when *V*_low_ was changed from -60mV to -70mV. We found that *f*_res_ decreased by 0.24±0.03 Hz and *Z*_max_ increased by 5.2±0.6 MΩ for high *ḡ_ca_* models, whereas *f*_res_ decreased by 0.07±0.02Hz and *Z*_max_ decreased by 2.6±0.2MΩ for low *ḡ_ca_* models (fig 9b, right panel). Therefore, neither model group reproduced the experimental changes in the Z-profile, specifically, a decrease in *f*_res_ and no change in *Z*_max_.

**Fig 9.**
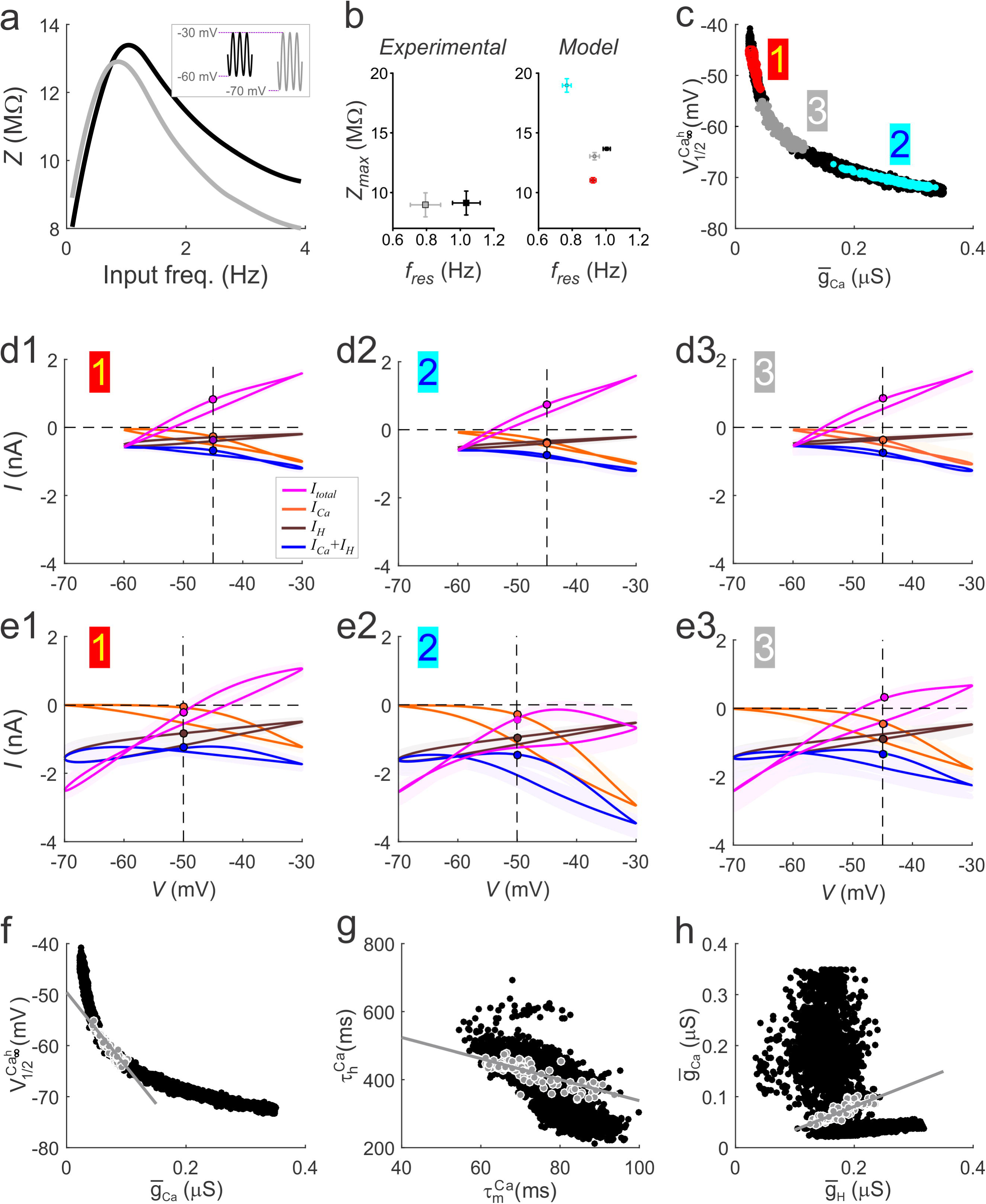
The effect of the lower voltage bound V_low_ of oscillations on *f*_res_ and *Z*_max_ constrains the optimal models. **a.** An example of the change in *Z*(*f*) measured in the biological PD neuron for V_low_ =-60mV (black line) and V_low_ =-70mV (grey line). Inset shows the bounds of voltage clamp inputs in the two cases**. b.** Shifting V_low_ from -60 mV to -70 mV lowers the value of *f*_res_ measured in the PD neuron significantly, without influencing Z_max_ (**b**. Experimental). i and Z_max_ values measured in a random subset of optimal model neurons corresponding to low or high *ḡ_ca_* values produced the same *f_re_*_s_ and Z_max_ values at *V*_low_ = -60mV (black dots), but distinct *f*_res_ and Z_max_ values at *V*_low_ = -70mV (low *ḡ_ca_*: red dots; high *ḡ_ca_*: cyan dots). A subset of optimal models could reproduce the experimental result in which *f*_res_ shifted to significantly lower values without affecting Z_max_. (grey dots). (**c)** *ḡ_ca_* - 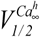 relationship separating out the different groups of models producing different responses to changes in *V*_low_ (colors correspond to **b** Model panel). Models depicted by grey dots are referred to as intermediate *ḡ_ca_* models. (**d1-e3**) mean voltage-gated ionic currents *I*_Ca_, *I*_H_ and *I*_Ca_+*I*_H_ and *I*_total_, shown as a function of voltage for V_low_ =-60 mV (**d1-d3**) and V_low_ = -70 mV (**e1-e3**). Numbers correspond to the location along the *ḡ_ca_* - 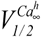 as shown in **c. f**. The intermediate *ḡ_ca_* models (grey dots) show a distinct 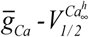 linear correlation. *g.* Intermediate *ḡ_ca_* models (grey dots) show a distinct and tighter 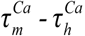 correlation compared to all optimal models (black dots). **h.** Intermediate *ḡ_ca_* models (grey dots) show a strong *ḡ_ca_* - *ḡ_H_* linear correlation that is not observed for all optimal models (black dots).

We consequently filtered the full optimal dataset (black dots fig 9c) to find a subset of models that reproduced the change in *f*_res_ and Z_max_ (to within 5% of the representative experimental Z(f) shown in fig 9a) when *V*_low_ was decreased to -70mV. Of the ^∼^9000 models in the population, we found ^∼^1000 models that produced the desired change. Interestingly, the resulting models showed a trade-off in values for *ḡ_ca_* and 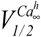 parameters that showed little overlap with the low and high *ḡ_ca_* model groups (fig 9c).

To understand why this particular group (which we will term intermediate *ḡ_ca_*) produced small changes in *Z_max_* when *V*_low_ was decreased, we plotted the current-voltage relationships for **I*_Ca_*, **I*_H_*, *I_Ca_*+*I_H_* and *I_total_* for V_low_ = -60 and - 70 mV, measured at f=1Hz (*f*_res_ at *V*_low_ = -60mV) and compared these models with the low and high *ḡ_ca_* models. For *V*_low_ =-60mV, the ionic currents behaved similarly for all model groups and *I_total_* was maximal at -30mV (magenta curve in fig 9d1-3), indicating the similarity of all models in the optimal dataset. However, when *V*_low_ was at -70mV revealed differences in peak *I*_Ca_, without affecting the peak amplitude of *I*_H_ across the different *ḡ_ca_* groups (fig 9e1-3). The differences in peak *I*_Ca_ accounted for most of the changes in *I*_total_ across the different *ḡ_ca_* groups. The *Z*_max_ values for intermediate *ḡ_ca_* models reproduced the small shift seen in experiments because *I*_Ca_ were at the correct level at high voltages (-30 mV) when *V*_low_ was at -70mV (fig 9e3). The other two groups did not produce appropriate Z_max_ for *V*_low_ = -70mV because either *I*_Ca_ was too small (and hence *I*_total_ too large), resulting in a smaller Z_max_ (fig 9e1) or vice versa (fig 9e2). It was also clear that the more negative voltages allowed for an increase in *I*_H_ levels and therefore larger contribution to the total current. With *V*_low_ at -70mV, not only was there a larger peak amplitude of *I*_H_ at the lower voltages, but the current at positive voltages also increased because of the very slow deactivation rate. Consequently, *I*_H_ did not fully turn off when *I*_Ca_ peaked, so that it also contributes to shaping the upper envelope of the total current. *I*_H_ kinetics were different across the groups (fig 9e1-e3). Taken together with the fact that when *I*_H_ was removed produced only parameter values with very high *ḡ_ca_* and very low 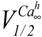 (fig S1), these data suggest that *I*_H_ could extend the range of I_Ca_ parameters over which MPR through compensation for variable levels of *I*_H_.

The *I_Ca_* in low *ḡ_ca_* models was too small when *V*_low_ was -70 mV, because the low conductance did not allow for a significant contribution from the additional de-inactivation (considering the higher 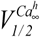 in this group) and therefore the peak current did not increase enough. Consequently, the contribution of *I_H_* at low voltages was greater than that of *I*_Ca_ at higher voltages (fig 9e2). Conversely, in the high *ḡ_ca_* group, 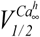 was more negative and so many more channels were available for de-inactivation and the contribution of *I*_Ca_ at higher voltages was much larger than that of *I*_H_ at low voltages (fig 9e3). These findings suggest that the balance between these two currents, that shape the lower and upper envelope of the total current response to voltage inputs, is necessary to produce the appropriate shift in *f*_res_ without influencing Z_max_ significantly.

The intermediate *ḡ_ca_* models were strongly correlated in *ḡ_ca_* - 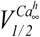 (R^2^ = 0.89; p < 0.001 fig 9f1, and had a stronger correlation in the 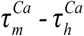 parameters compared to all models (R^2^ = 0.65; p < 0.001; fig 9g). Limiting the optimal models to the intermediate *ḡ_ca_* group also revealed a correlation in the *ḡ_ca_* - *ḡ_H_* parameters (R^2^ = 0.79; p < 0.001; fig 9h). This new correlation may be produced by the balance of the amplitudes of *I*_H_ and *I*_Ca_ at the lower and higher voltages, respectively.

#### *f_res_* and *Q_z_* are maintained by distinct pairwise correlations

To determine if any of the MPR attributes were sensitive to the correlations, we ran a 2D sensitivity analysis on a random subset of 50 models. We tested for significant difference in sensitivity across low, intermediate and high levels of *ḡ_ca_*. In particular, we tested for significant sensitivity of *f*_res_ *and Q_Z_* when parameters were co-varied in directions parallel (L) or perpendicular (L^⊥^) to their respective population correlation lines.

We first examined whether *f*_res_ and *Q_Z_* were sensitive to 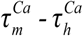 for both high (fig 10a1), low (fig 10a2), and intermediate *ḡ_ca_* (fig 10a3) when parameters were moved along L and L^⊥^ (blue and green line; fig 10a1-a3). For high and intermediate *ḡ_ca_* models, *f*_res_ sensitivities in the L group were negative and not significantly different (3-way RM ANOVA; N=50, p > 0.05), but both groups were significantly different from the low *ḡ_ca_* group (3-way RM ANOVA; N=50, p < 0.001), which had a positive sensitivity (fig 10b). This result indicates that the correlation did a better job at maintaining the value of *f*_res_ when the value of *ḡ_ca_* is intermediate or high. For all *ḡ_ca_* groups, we found that there was a significant interaction between the Z attribute and direction (2-way RM ANOVA; F(1, 49) = 853.52, p < 0.001). When carrying out a pairwise comparison for, we found a significant difference in sensitivity between L and L^⊥^ for f*_res_* (t(93.57)=28.251, p<0.001). Similarly, for all *ḡ_ca_* groups, significant difference in sensitivity between L and L^⊥^ for Q_Z_ (t(93.57)=-8.294, p<0.001). Because the difference between L and L^⊥^ for Q_Z_ was negative, these results suggest that the 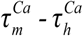 correlation determines *f*_res_ and not Q_Z_ (fig 10b).

**Fig 10.**
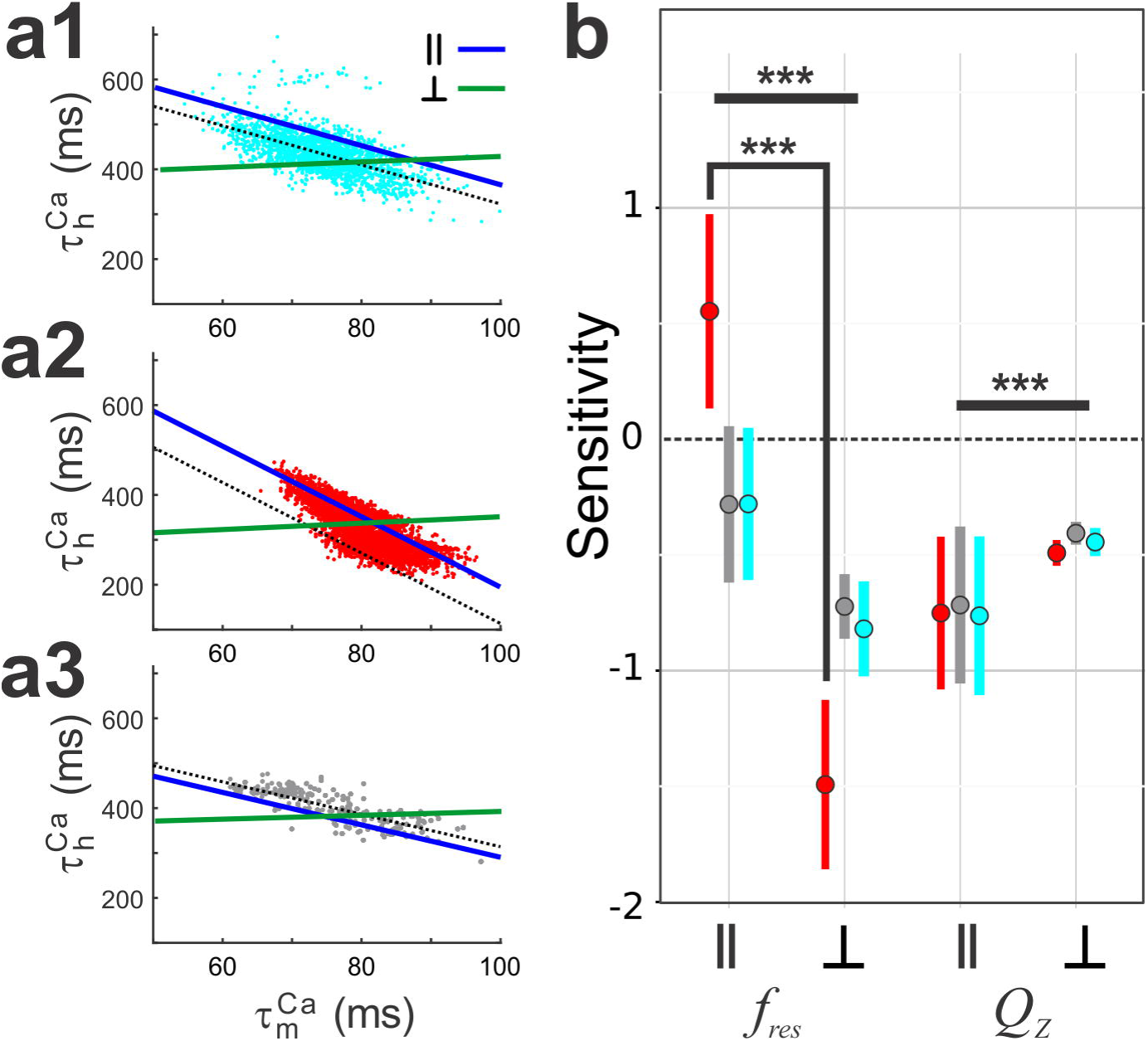
Assessing the dependence of *f*_res_ and *Q_Z_* on the 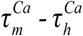 linear correlation. **a**. Parameter values for each model were changed along a line parallel (, blue) to the correlation line (black) or along a perpendicular line (^⊥^ grey). This was done for models with high (cyan; **a1**), low (red; **a2**) and intermediate (grey; **a3**) *ḡ_ca_* models. For each model and each line, or ^⊥^, we fit a line to the relative change in either *f*_res_ or Q_Z_ as a function of the relative change in **ḡ_ca_**. **b.** The sensitivity values of *f*_res_ or *Q_Z_* to or ^⊥^ are shown for the three groups. **c.** Impedance profiles showing how Q_Z_ changes when the parameters vary along a line parallel (blue) or perpendicular (grey) to the *ḡ_Ca_* - *ḡ_H_* correlation line in one optimal model. Arrows show the direction of the movement of Z_max_ and *f*_res_ for the change in parameters along or ^⊥^ for the high (**c1**), low (**c2)** and intermediate (**c3**) **ḡ_ca_** model.

We next examined whether *f*_res_ and *Q_Z_* were sensitive to the *ḡ_ca_* - 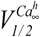 correlation for the three model groups (fig 11a1-3). For all *ḡ_ca_* groups, we found that there was a significant interaction between the Z attribute and direction (2-way RM ANOVA; F(1, 49) = 1262.73.2, p < 0.001). When carrying out a pairwise comparison for each direction within an attribute, we found a significant difference in sensitivity between L and L^⊥^ for f*_res_* (t(95.18)=10.10, p<0.001). Similarly, for all *ḡ_ca_* groups, we found a significant difference in sensitivity between L and L^⊥^ for Q_Z_ (t(95.18)=-35.62, p<0.001). Therefore, these results suggest that the *ḡ_ca_* - 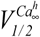 correlation determines Q_Z_ and not *f*_res_ (fig 11b).

**Fig 11.**
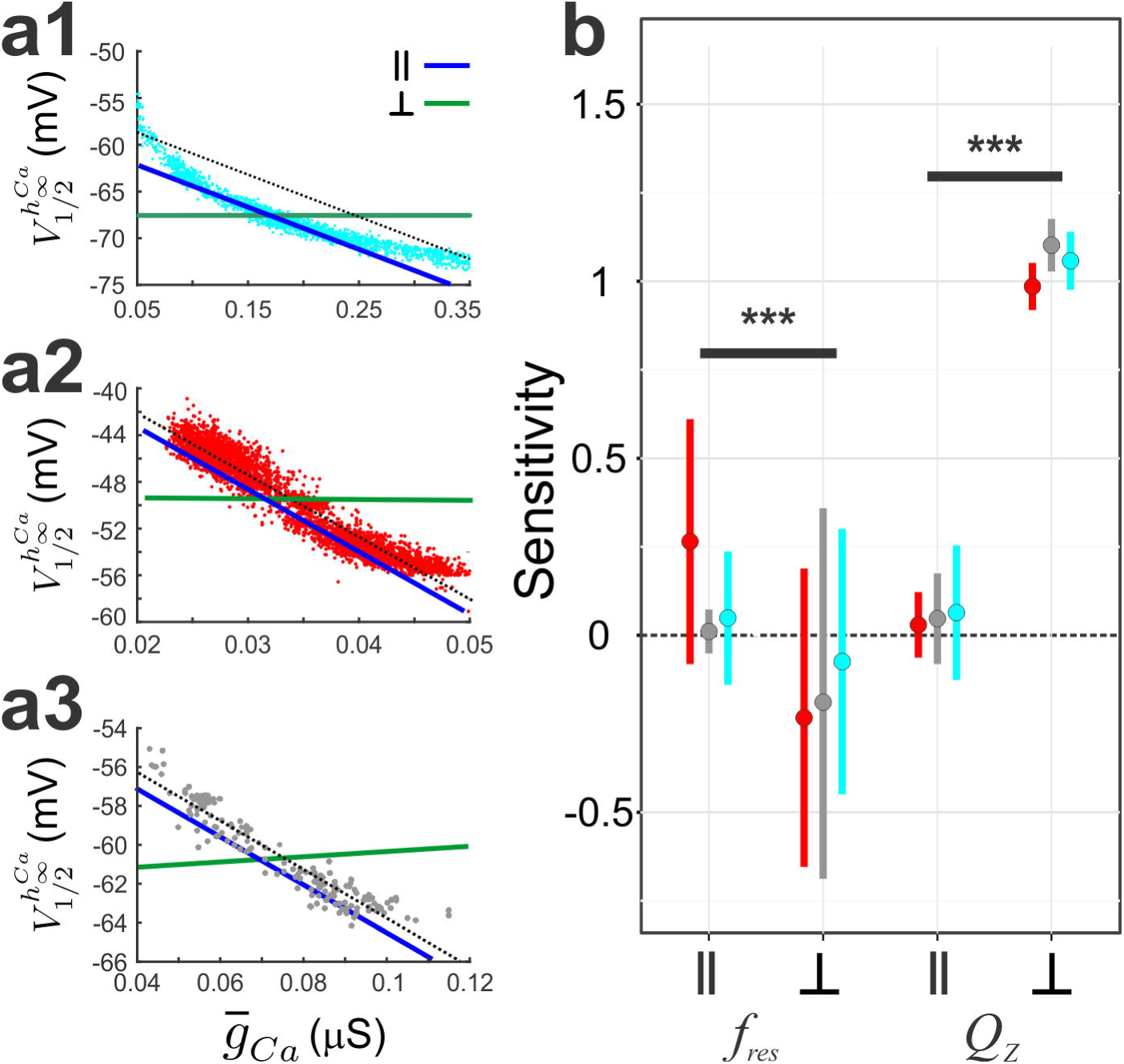
Assessing the dependence of *f_res_* and *Q_Z_* on the linear 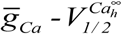 correlation. **a.** Parameter values for each model were changed along a line parallel (, blue) to the correlation line (black) or along a perpendicular line ^⊥^grey). This was done for models with high (cyan; **a1**), low (red; **a2**) and intermediate (grey; **a3**) *ḡ_Ca_* models. For each model and each line, or ^⊥^, we fit a line to the relative change in either *f*_res_ or Q_Z_ as a function of the relative change in *ḡ_Ca_*. **b.** The sensitivity values of *f*_res_ or *Q_Z_* to or ^⊥^ are shown for the three groups. **c.** Impedance profiles showing how Q_Z_ changes when the parameters vary along a line parallel (blue) or perpendicular (grey) to the *ḡ_Ca_* - 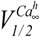 correlation line in one optimal model. Arrows show the direction of the movement of Z_max_ and *f*_res_ for the change in parameters along or ^⊥^ for the high (**c1**), low (**c2**) and intermediate (**c3**) *ḡ_Ca_* model.

Finally, we tested the sensitivity of *f*_res_ and Q_Z_ to the *ḡ_ca_* - *ḡ_H_* correlation in the intermediate *ḡ_ca_* group (fig 12a). We found that there was a significant interaction between the Z attribute and direction (2-way RM ANOVA; F(1, 11.12) = 2236.2, p < 0.001). When carrying out pairwise comparisons between directions for each attribute, we found there was a significant difference in *f*_res_ sensitivity between L and L^⊥^ (t(93.93) = 2.65, p = 0.0095; fig 12). Although the sensitivity of Q_Z_ was not 0 for L, the difference in sensitivity values between L and L^⊥^ was also significantly different (t(93.93) = 62.157, p < 0.0001; fig 12b). These results suggest that, when *V*_low_ is at -70 mV, for this subset of models to shift *f_re_*_s_ with only small shifts in Z_max_, *ḡ_H_* and *ḡ_ca_* values must be balanced. It may be possible that the Q_Z_ sensitivity is not closer to zero along L because 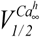, which is also negatively correlated with *ḡ_ca_*, should decrease too to compensate for changes in Q_Z_.

**Fig 12.**
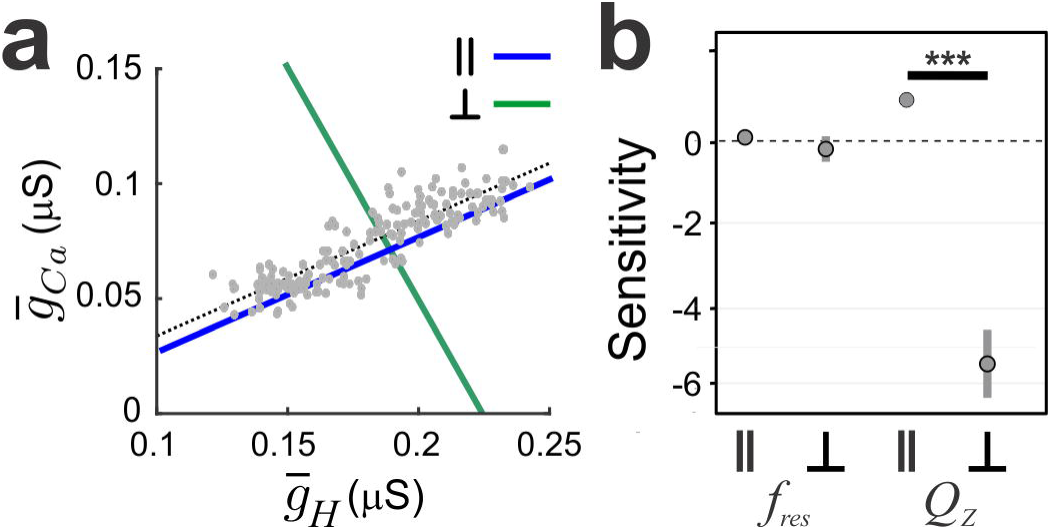
Assessing the dependence of ***f*_res_** and ***Q_Z_*** of the intermediate ***ḡ_Ca_*** models on the linear ***ḡ_Ca_ - *ḡ_H_**** correlation. **a**. Parameter values for each model were in the intermediate *ḡ_Ca_* group (see fig 9) were changed along a line parallel (, blue) to the correlation line (black) or along a perpendicular line (^⊥^ grey). For each model and each line, or *-* ^⊥^ we fit a line to the relative change in either *f*_res_ or *Q_Z_* as a function of the relative change in *ḡ_Ca_***. b.** The sensitivity values of *f*_res_ or *Q_Z_* to or ^⊥^ are shown for the three groups. **c.** Impedance profiles showing how *Q_Z_* changes when the parameters vary along a line parallel (blue) or perpendicular (grey) to the *ḡ_Ca_ - *ḡ_H_** correlation line in one optimal model. Arrows show the direction of the movement of Z_max_ and *f*_res_ for the change in parameters along or ^⊥^.

## Discussion

Many neuron types exhibit membrane potential resonance (MPR) in response to oscillatory inputs. Several studies have shown that the resonant frequency of individual neurons is correlated with the frequency of the network in which they are embedded [2, 6, 12, 14, 22, 46]. Moreover, networks of resonant neurons have been proposed to generate more robust network oscillations than neurons with low-pass filter properties [27, 28]. In several cases, the underlying nonlinearities and time scales that shape the Z-profile also shape specific properties of the spiking activity patterns, thus leading to a link between the subthreshold and suprathreshold voltage responses [25, 47].

Previous work in the crustacean stomatogastric pyloric network has shown that the resonance frequency of the pyloric pacemaker PD neurons is correlated with the pyloric network frequency and is sensitive to blockers of both *I*_H_ and *I*_Ca_ [12-14]. However, it was not clear how these voltage-gated ionic currents and the passive properties could interact to generate MPR in the PD neurons. Previous modeling work showed that these currents participate in the generation of resonance in CA1 pyramidal neurons [16, 17]. However, due to the differences in *I*_Ca_ time constants, the interaction between its activating and inactivating gating variables did not produce phasonance in CA1 pyramidal neurons, while it does in PD neurons. On a more general level, it is not well understood how the nonlinear properties of ionic currents affect their interplay. Previous studies have shown such interactions may lead to unexpected results, which are not captured by the corresponding linearizations [16-19]. This complexity is expected to increase when two currents with resonant components are involved [16, 48]. We therefore set out to investigate the biophysical mechanism underlying such interactions by using a combined experimental and computational approach and the biological PD neuron as a case study. The two PD neurons are electrically coupled to the pacemaker anterior burster neuron in the pyloric network and their MPR directly influences the network frequency through this electrical coupling [22]. Consequently, our findings have a direct bearing on how the pyloric network frequency is controlled.

Many studies of biophysical models have explored the parameter space using a brute-force technique, by sampling the parameters on a grid [40, 49]. Although this technique provides a rather exhaustive sampling of the parameter space, using a fine grid on a large number of free parameters could lead to combinatorial explosion and result in a prohibitive number of simulations. On the other hand, a sparse sampling may miss “good” solutions. A multi-objective evolutionary algorithm (MOEA) can generate multiple trade-off solutions in a single run and can handle large parameter spaces very well. In contrast to a brute-force approach, the MOEA can potentially cover a much larger range with possibly hundreds of values [38]. One disadvantage of the MOEA is that, as the number of objectives increases, the search may miss a large portion of the parameter space. This occurs because randomly generated members often tend to be just as good as others, which means that the MOEA would run out of room to introduce new solutions in a given generation. To try to overcome this problem, we carefully chose the parameters of the MOEA such as population size, mutation and crossover distribution indices (100, 20 and 20, respectively) and ensured that the sampled population covered the parameter space evenly. Additionally, we ran the MOEA multiple times, each time collecting all the good parameter sets until one has exhausted all regions of the parameter space where good models exist.

In previous work, we and other authors have examined how the additive interaction of ionic currents with resonant and amplifying gating variables shape the Z and ϕ profiles at both the linear and nonlinear levels of description [6, 15, 18, 20, 32, 33, 50]. However, the role of inactivating currents in the generation of MPR is not so clear. Authors have established that *I*_Ca_ can generate MPR in the absence of additional ionic currents [21], that the activation variable diminishes the propensity for MPR and the interaction with *I*_H_ enhances the dynamic range of parameters producing *I*_Ca_-mediated resonance [16]. Even so, to date, only a descriptive explanation of how the ionic current parameters affect certain attributes of MPR has been provided, but no study has provided a mechanistic understanding in terms of the parameters of *I*_Ca_ that go beyond numerical simulations.

Similar to [16], the model we used in this paper involves the interaction between resonant and amplifying components. Specifically, this model includes a calcium current with both activation (amplifying) and inactivation (resonant) gating variables, and an H-current with a single activation (resonant) gate. Since *I*_H_ and *I*_Ca_ shape the lower and upper envelopes of the voltage response to current inputs, respectively [12], given the appropriate voltage-dependence and kinetics of the currents both could play equal roles at different voltage ranges. In fact, either *I*_Ca_ inactivation or *I*_H_ is capable of producing MPR [2, 21]. In CA1 pyramidal neurons, the differences in Z profiles are due to the passive properties and the kinetics of *I*_H_ [4]. It is possible that the kinetic parameters of *I*_H_ and *I*_Ca_ are tuned so that they contribute nearly equally to shaping the envelopes of the voltage-clamp current.

By tracking the current response to sinusoidal voltage inputs at various frequencies, we found that the *f*_res_ and *f*_ϕ=0_ are driven by the peak phase of *I*_Ca_, and that *f*_res_ and *f*_ϕ=0_ are nearly equal because of the phase matching of *I*_Ca_ with *I*_L_. This is not always the case for neuronal models, and dynamical systems in general, not even for linear models, except for the harmonic oscillator [18-20]. In fact, as we mentioned above, this is not the case for the *I_Ca_* model used in [16], although our results on the *I*_Ca_ inactivation time constant are consistent with that study. In these models phase advance for low input frequencies required the presence of **I*_H_.* The underlying mechanisms are still under investigation and are beyond the scope of this paper. However, the fact that it occurs was crucial to develop a method to investigate the dependence of the resonant properties, particularly the dependence of the *f*_res_ on the *I_Ca_* time constants, using phase information. To date, no other analytical method is available to understand the mechanisms underlying this type of phenomenon in voltage clamp. The tools we developed are applicable to other neuron types for which *f*_res_ is equal to or has a functional relationship with *f*_φ=0_. However, the conditions under which such a functional relationship exists still needs to be investigated.

Linear correlations between biophysical parameters of the same or different currents have been reported [37] and may be important in preserving the activity of the model neuron and its subthreshold impedance profile attributes [41]. Previous studies examined combinations of parameters in populations of multi-compartment conductance-based models fit to electrophysiological data [16, 51] and found only weak pairwise correlations suggesting that the correlations do not arise from electrophysiological constraints. In contrast, constraining the parameters of the ionic currents found to be essential for MPR in PD neuron by MPR attributes, we observed strong correlations underlying parameters when the *Z* and φ were constrained by the experimental data. We found that constraining the model parameters by *f*_res_ produced a correlation between the values of time constants of *I*_Ca_ among the population of ^∼^9000 optimal parameter sets. Furthermore, running a 2D sensitivity analysis confirmed that the time constants were constrained so that the effect of making inactivation slower was compensated for by making activation faster to maintain *f*_res_ constant.

The optimal model parameter sets showed a nonlinear co-variation relationship between the *ḡ_ca_* and half-inactivation voltage of *I*_Ca_. However, the models could be divided into two groups, low and high *ḡ_ca_* in each of which this co-variation was close to linear. Interestingly, although *I*_Ca_ alone was the primary current underlying MPR, in the absence of *I*_H_ (with *ḡ_H_* = 0) the models were restricted to the high *ḡ_ca_* group. A 2D sensitivity analysis showed that co-varying parameters in each groups along their respective correlation lines preserved Q_Z_ without affecting *f*_res_, indicating that each group requires a distinct changes in one parameter to compensate for effects of changes in the other. Local sensitivity analysis showed that changes in 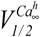 had opposite effects on *f*_res_ between high and low *ḡ_ca_* groups. Increasing 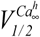 decreased *f*_res_ in high *ḡ_ca_* models but increased it in low *ḡ_ca_* models. A previous modeling study has found that changes in 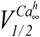 greatly influenced the amplitude of MPR with little effect on post-inhibitory rebound in thalamic neurons [21]. It would be interesting to verify whether the mechanisms that generate MPR overlap with those that contribute to post-inhibitory rebound properties.

Previous work in our lab has shown that the voltage range of oscillations significantly affects *f*_res_ [13]. Here we show that decreasing, *V*_low_, the lower bound of the oscillation voltage of the PD neuron, from - 60 to -70 mV, significantly shifted *f*_res_ to smaller values without affecting Z_max_. Within our optimal model parameter sets, we obtained a set of ^∼^1000 models in the intermediate *ḡ_ca_* range that produced a similar shift in *f*_res_ but no change in Z_max_. Because *V*_low_ greatly affects both *I*_Ca_ inactivation and *I*_H_ activation, this indicated a potential interaction between these two currents. In fact, we found that because *I*_H_ and *I*_Ca_ are activated preferentially in different voltage ranges, their amplitudes needed to be balanced to keep Z_max_ unchanged when *V*_low_ was decreased. If the ratio of *I*_H_ to *I*_Ca_ amplitudes is incorrect, then *Z* will amplify (for high *ḡ_ca_* models) or attenuate (for low *ḡ_ca_* models). The intermediate *ḡ_ca_* models also showed a stronger 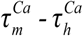 correlation, which may be important in matching the phase of *I*_Ca_ with that of *I*_L_. This group also showed a strong *ḡ_H_* − *ḡ_ca_* correlation, which may provide a mechanism for controlling the changes in *I*_H_ amplitude at more negative voltage with similar changes in *I*_Ca_ amplitude at more positive voltages.

In contrast to the findings of Rathour and Narayanan [16], in our optimal models the *I*_H_ amplitude was not different across the groups with different *I*_Ca_ properties. However, since *I*_Ca_ and *I*_H_ are differentially modulated [45, 52], their functional role may overlap when their voltage thresholds and time constants are shifted by neuromodulation. Therefore, we expect that under certain neuromodulatory contexts, *I*_H_ may play more of an active role in the generation of MPR. A similar effect of two ionic currents on resonance has been observed in the hippocampal pyramidal cells that participate in the theta rhythm, in which two currents, the slow potassium M-current and *I*_H_, were found to operate at the depolarized and hyperpolarized membrane potentials respectively to generate theta-resonance [2].

In general, variability of ionic current expression in any specific neuron type should lead to great variability in network output. Yet, network output in general, and specifically the output of the crustacean pyloric network is remarkably stable across animals [30, 53, 54]. Our results suggest that in oscillatory networks the interaction among ionic currents in an individual neuron may be tuned in a way that the variability of the output is reduced in response to oscillatory inputs. Although our computational study may provide some insight into how such stability is achieved, it also indicates a need for additional mathematical analysis to elucidate the underlying mechanisms.

## Methods

### Electrophysiology

The stomatogastric nervous system of adult male crabs (*Cancer borealis*) was dissected using standard protocols as in previous studies [14]. After dissection, the entire nervous system including the commissural ganglia, the esophageal ganglion, the stomatogastric ganglion (STG) and the nerves connecting these ganglia, and motor nerves were pinned down in a 100mm Petri dish coated with clear silicone gel, Sylgard 186 (Dow Corning). The STG was desheathed to expose the PD neurons for impalement. During the experiment, the dish was perfused with fresh crab saline maintained at 10487 13°C. After impalement with sharp electrodes, the PD neuron was identified by matching intracellular voltage activity with extracellular action potentials on the motor nerves. After identifying the PD neuron with the first electrode, a second electrode was used to impale the same neuron in preparation for two-electrode voltage clamp. Voltage clamp experiments were done in the presence of 10^-7^ M tetrodotoxin (TTX; Biotium) superfusion to remove the neuromodulatory inputs from central projection neurons (decentralization) and to stop spiking activity [13, 14].

Intracellular electrodes were prepared by using the Flaming-Brown micropipette puller (P97; Sutter Instruments) and filled with 0.6M K_2_SO_4_ and 0.02M KCl. For the microelectrode used for current injection and voltage recording, the resistance was, respectively, 10-15MΩ and 25-35MΩ. Extracellular recording from the motor nerves was carried out using a differential AC amplifier model 1700 (A-M Systems) and intracellular recordings were done with an Axoclamp 2B amplifier (Molecular Devices).

### Measuring the Z-profile

During their ongoing activity, the PD neurons produce bursting oscillations with a frequency of ^∼^1 Hz and slow-wave activity in the range of -60 to -30 mV. Activity in the PD neuron is abolished by decentralization. The decentralized PD neuron shows MPR in response to ZAP current injection when the current drives the PD membrane voltage to oscillate between -60mV and -30mV, which is similar to the slow-wave oscillation amplitude during ongoing activity [12]. The MPR profiles are not significantly different when measured in current clamp and voltage clamp [14]. Since the MPR depends on the dynamics of voltage-gated ionic currents, it will also depend on the range and shape of the voltage oscillation. Therefore, to examine how *Z*(*f*) in a given voltage range constrains the properties of voltage-gated currents and how factors that affect the voltage range change MPR, we measured *Z*(*f*) in voltage clamp [10].

To measure the Z-profile, the PD neuron was voltage clamped with a sweeping-frequency sinusoidal impedance amplitude profile (ZAP) function [55] and the injected current was measured [14]. To increase the sampling duration of lower frequencies as compared to the larger ones, a logarithmic ZAP function was used:

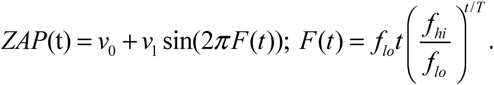

The amplitude of the ZAP function was adjusted to range between -60 and -30 mV (*v*_0_=-45 mV, *v*_1_=15 mV) and the waveform ranged through frequencies of *f*_lo_=0.1 to *f_hi_*=4 Hz over a total duration T=100 s. Each ZAP waveform was preceded by three cycles of sinusoidal input at *f_lo_* which smoothly transitioned into the ZAP waveform. The total waveform duration was therefore 130 s.

Impedance is a complex number consisting of amplitude and phase. To measure impedance amplitude, we calculated the ratio of the voltage and current amplitudes as a function of frequency and henceforth impedance amplitude will be referred to as *Z*(*f*)). To measure *φ_z_*(*f*), we measured the time difference between the peaks of the voltage clamp ZAP and the measured clamp current. One can also measure *Z*(*f*) by taking the ratio of the Fourier transforms of voltage and current. However, spectral leakage, caused by taking the FFT of the ZAP function and the nonlinear response, often resulted in a low signal-to-noise ratio and therefore in inaccurate estimates of impedance. Such cases would lead to less accurate polynomial fits compared to the cycle-to-cycle method described above and we therefore limited our analysis to the cycle-to-cycle method.

Because the average Z-profile may not be a realistic representation of a biological neuron, we used the attributes of *Z* and φ measurements from a single PD neuron as our target. We characterized attributes of *Z* into five objective functions used for fitting by specifying five points of the profile (fig 1a). These five points were:

- (*f*_0_, *Z*_0_), where *Z*_0_ = *Z*(*f*_0_) and *f*_0_ = 0.1 Hz,
- (*f_res_*, *Z*_max_), thereby capturing *Qz* = *Z*_max_ - *Z*_0_,
- (*f*_1_, *Z*(*f*_1_)) where *f*_1_ = 4 Hz,
- The two frequencies at which *Z* = *Z*_0_ + *Q_Z_* / 2. Pinning the profile to these points captures the frequency bandwidth Λ ½ which is the frequency range for which *f* > *Z*_0_ + *Q_Z_*/ 2 (fig 1a).

We also constructed five objective functions to capture the attributes of φ(*f*) at five points (fig 1b):

- (*f*_0_, *φ*(*f*)),
- (*f_φ_*=_0_, 0), where *f_φ_*,_=0_, is the phasonant frequency
- (*f_φmax_*, *φ*_max_) where *φ*_max_ is the maximum phase advance,
- (*f_φmin_*, *φ*_min_) where *φ*_min_ is the maximum phase delay,
- (2 Hz, *φ_f_*_=_*_2_*) capturing the phase at 2Hz.

### Single-compartment model

We used a single-compartment biophysical conductance-based model containing only those currents implicated in shaping *Z* and *φ* [12]. We performed simulations in voltage clamp and measured the current as:

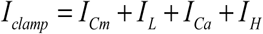
where *I*_Cm_ is the capacitive current 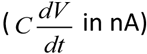, C_m_ is set to 1 nF and *I*_L_ is the voltage-independent leak current in nA. The voltage-dependent currents *I*_curr_ (*I*_Ca_ or *I*_H_) in nA are given by

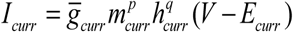
where *V* is the ZAP voltage input (see below), *m_curr_* is the activation gating variable, *h_curr_* is the inactivation gating variable, 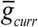 is the maximal conductance in μS, *E_curr_* is the reversal potential in mV, and *p* and *q* are non-negative integers. For *I_Ca_*, *p* = 3, *q* = 1 and, for *I_H_*, *p* = 1 and *q* = 0. The generic equation that governs the dynamics of the gating variables is:

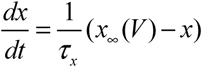

Where *x* = *m*_curr_ or *h*_curr_, and

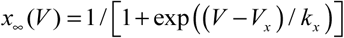

The sign of the slope factor (*k_x_*) determines whether the sigmoid is an increasing (negative) or decreasing (positive) function of *V*, and *V_x_* is the midpoint of the sigmoid.

A total of 8 free model parameters were defined (Table 1), which were optimized in light of the objective functions introduced above, to yield a good fit to the Z-profile attributes as described below.

The slope factors *k_x_* of the sigmoid functions 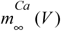,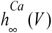 and 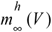 were fixed at -8 mV, 6 mV, and -7 mV, respectively. 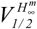 was fixed at -70 mV, using data from experimental measurements in crab [44]. The voltage-dependent time constant for *I_H_* was also taken from [44] to be

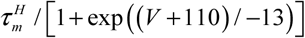
where the range of 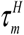 is given in Table 1.

### Fitting models to experimental Data

Computational neuroscience optimization problems have used a number of methods, such as the “brute-force” exploration of the parameter space [51] and genetic algorithms [56]. However, the brute-force method is computationally prohibitive for an 8-dimensional model parameter space, which would require potentially very fine sampling to find optimal models. [57]. We used an MOEA (evolutionary optimization) to identify optimal sets of model parameters constrained by experimental *Z* and φ attributes. MOEAs are computationally efficient at handling high-dimensional parameter spaces and other studies have used them to search for parameters constrained by other types of electrophysiological activity [57]

Evolutionary optimization finds solutions by minimizing a set of functions called objective functions, or simply objectives, subject to certain constraints. In our problem, each objective represents the Euclidean distance between the target and the model attributes of *Z* and *φ*. When optimizing multiple (potentially conflicting) objectives, MOEA will find a set of solutions that constitute trade-offs in objective scores. For instance, an optimal parameter set may include solutions that are optimal in *f*_res_ but not in Q_z_ or vice versa and a range of solutions in between that result from the trade-offs in both objectives. In this paper, we used the non-dominated sorting genetic algorithm II (NSGA-II) [38, 58] to find optimal solutions, which utilizes concepts of non-dominance and elitism, shown to be critical in solving multi-objective optimization problems [58]. Solution x_1_ is said to dominate solution x_2_ if it is closer to the target *Z*(*f*) and φ(*f*) profiles in at least one attribute (e.g., *f*_res_) and is no worse in any other attributes (e.g., *Q_Z_*, *Z*_0_, etc.).

NSGA-II begins with a population of 100 parameter combinations created at random within pre-determined lower and upper limits (Table 1). The objective values for each parameter combination are calculated and ordered according to dominance. First, the highest rank is assigned to all of the non-dominated, trade-off solutions. From the remaining set of parameters, NSGA-II selects the second set of trade-off solutions. This process continues until there are no more parameter combinations to rank. Genetic operators such as binary tournament selection, crossover, and mutation form a child population. A combination of the parent and child parameter sets form the population used in the next generation of NSGA-II [38, 58]. NSGA-II favors those parameter combinations—among solutions non-dominating with respect to one another—that come from less crowded parts of the parameter search space (i.e., with fewer similar, in the sense of fitness function values, solutions), thus increasing the diversity of the population. The crowding distance metric is used to promote large spread in the solution space [38].

We ran NSGA-II multiple times (3-5 times, until the mean values of the distributions of optimal parameters was stable) each time for 200 generations with a population size of 100, and pooled the solutions at the end of each run to form a combined population of ^∼^9000 parameter combinations. The algorithm stopped when no additional distinct parameter combinations were found. The *Z* and *φ* values associated with the optimal parameter sets match the target features (objectives) defining *Z* and *φ* to within 5% accuracy.

To test whether two parameters were significantly correlated in the population of 9000 PD models, we calculated the Pearson’s correlation coefficients for each pair of parameters and used a permutation test to determine the number of times the calculated correlation coefficient (using a random subset of 20 models). The p-value was given as the fraction of R-values for the permuted vectors greater than the R-value for the original data [51]. We also used a t-test to determine whether the calculated slope of the linear fit differed significantly from zero, which gave us identical results. We repeated both procedures 2000 times, each time with a random subset of 20 models and calculated the percentage of times we obtained a p-value < 0.01.

### Sensitivity Analysis

We assessed how the values of *f*_res_ and *Q_Z_* depend on changes in parameter values by performing a sensitivity analysis as in [59]. We split the model parameters into two categories: additive, for the voltage-midpoints of activation and inactivation functions, and multiplicative, for the maximal conductances and time constants. We changed the parameters one at a time and fit the relative change in the resonance attributes as a linear function of the relative parameter change. We changed the multiplicative parameters on a logarithmic scale to characterize parameters with both low and high sensitivity.

Multiplicative parameters were varied as *p*_n+1_ = exp(±Δ*p*_n_) *p*_0_ with *Δρ_η_* = 0.001*1.15^n^ and the sign indicating whether the parameter was increased or decreased. To ensure approximate linearity, we added points to the fit until the *R^2^* value fell below 0.98. The sensitivity was defined as the slope of this linear fit (fig 2). For example, if a resonance attribute has a sensitivity of 1 to a parameter, then a 2-fold change in the parameter results in a 2-fold change in the attribute. We changed additive parameters by ±0.5 mV.

We assessed the sensitivity of *f*_res_ and *Q*_z_ to parameter pairs (p_1_ and p_2_) that were correlated. We first fit a line through the correlated values in the p_1_-p_2_ space. We then shifted this line to pass through a subset of 50 random points in p_1_-p_2_ space, resulting in a family of parallel lines, L. For each point, we also produced a line perpendicular to a line L^⊥^. For each model, we performed a sensitivity analysis as before but used the linear fit equation L or L^⊥^ to calculate value of p_2_. We fit the relative change in the Z(f) attribute as a linear function of the correlated change in p_1_ and p_2_. We used the slope of the linear fit to represent the sensitivity. We used a 2-and 3-way repeated measures ANOVA and the lsmeans function in R to perform pairwise comparisons of means in testing for significant differences between each group of g_Ca_, each direction, L and L^⊥^ and between each *Z* attribute, *f*_res_ and Q_Z_.

For each model, we solved a system of three differential equations for *m*_H_, *m*_Ca_ and *h*_Ca_ (voltage was clamped). All simulations were performed using the modified Euler method [60] with a time step of 0.2 ms. The simulation code, impedance calculations, and MOEA were written in C++. MATLAB (The MathWorks) and R were used to perform statistical analyses.

## Supporting Information Legends

**S1. Changing the value of 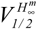 does not change the correlations observed among the model parameters. a.** Correlations shown in Fig. 8b with 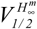 at -70 mV. **b**. Correlations obtained with 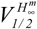 set to -96 mV (red dots). MOEA was run only once in this case, compared to 5 times in panel a (hence the difference in the number of points). Black dots are the same as panel **a.** Note that the values of *ḡ_H_* in this case are about 10 times larger than those in panel **a,** but the correlations (green boxes) remain intact. More importantly, the range of parameters other than *ḡ_H_* is exactly the same in both cases.

**S2. *I*_H_ extends the dynamic range of *I*_Ca_ parameters over which *I*_Ca_-mediated MPR occurs.** Parameter values for the optimal models in *ḡ_ca_* - 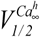 space shown for all models (grey dots) and those without *I*_H_ (blue dots). We removed I_H_ by setting *ḡ_H_* = 0, and ran the MOEA multiple times using the same Z- and *φ*-profiles to constrain the I_Ca_ parameters. A linear fit (green) shows that, when *ḡ_H_* =0, the relationship between *ḡ_ca_* - 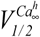 is linear and matches a narrow range of the high *ḡ_ca_* values in fig 8c.

